# A pesticide hijacks a neurohormonal circuit to block ovulation in insects

**DOI:** 10.64898/2026.06.30.735649

**Authors:** Ji-Yang Xing, Shao-Cong Su, Pin-Xuan Lin, Yue Guo, De-Run Wu, Ren-Ai Li, Dick R. Nässel, Chris Bass, Jie Chen, Cong-Fen Gao, Shun-Fan Wu

## Abstract

The global decline of insect populations threatens ecosystem stability, yet the hidden physiological impacts of sublethal pesticide exposure remain largely uncharted. Here we show that emamectin benzoate (EB), a widely used insecticide, suppresses female fertility not by disrupting egg development, but by specifically blocking ovulation. Through a systematic genetic screen in Drosophila, we identify the glutamate-gated chloride channel GluClα in a discrete set of octopaminergic neurons as the primary neural target. EB subverts the activity of this circuit, thereby impairing octopamine-dependent follicle rupture, suppressing ovarian ecdysteroid signaling, and altering oviduct muscle dynamics—three coordinated processes essential for egg release. Strikingly, this ovulatory blockade is recapitulated in diverse dipteran pests and disease-vector mosquitoes, revealing a conserved vulnerability. Our findings establish a neurohormonal mechanism by which a common agrochemical hijacks reproductive control, and they expose ovulation as a critical, hitherto unrecognized nexus between environmental chemicals and insect population dynamics.

**Significance:** Sublethal pesticide exposure can profoundly alter insect physiology beyond acute toxicity, yet the underlying mechanisms remain unclear. Here, we show that emamectin benzoate (EB), a widely used avermectin insecticide, suppresses ovulation and female fecundity in *Drosophila* by disrupting octopaminergic signaling, follicle rupture, and oviduct muscle contraction. Mechanistically, EB acts through GluClα-dependent neural pathways to impair ovulation-related processes and reproductive muscle dynamics. Notably, these inhibitory effects are conserved across multiple dipteran species, suggesting that reproductive suppression represents a broader physiological consequence of EB exposure. Our findings reveal a previously unrecognized neuro-reproductive mechanism of pesticide action and provide new insight into how sublethal insecticide exposure may influence insect population dynamics.

## INTRODUCTION

Global insect declines have become a major ecological concern, with habitat loss, climate change, pollution, and the widespread use of insecticides recognized as important anthropogenic drivers (1–3). Because reproduction is a fundamental determinant of population growth and persistence, chemical interference with reproductive physiology may represent an underappreciated mechanism by which environmental stressors reshape insect population dynamics (4–7). In natural and agricultural environments, insects are frequently exposed to sublethal concentrations of insecticides that do not cause immediate lethality but can alter development, behavior, reproduction, and life span (1, 3, 8). However, these effects are often interpreted as indirect consequences of generalized chemical stress, and the mechanisms by which sublethal insecticide exposure disrupts specific physiological systems remain poorly understood (9, 10). A central challenge in environmental toxicology is thus to determine whether sublethal pesticide exposure simply causes general physiological impairment or rather perturbs specific regulatory systems that control key biological processes (1, 3, 12).

Many insecticides act on ion channels in the nervous system, raising the possibility that their effects may extend beyond acute toxicity to influence neural regulation of physiology (13–17). In insects, reproductive processes are tightly controlled by neurohormonal pathways (18, 19). In *Drosophila melanogaster*, ovulation is governed by coordinated neurohormonal and physiological processes (20–22), involving octopaminergic signaling (23, 24), hormonal regulation (25), and follicle rupture (26, 27). Disruption of this integrated system leads to egg retention and reduced fertility (25, 26). However, whether sublethal chemical exposure perturbs such coordinated regulatory networks to influence ovulation remains unknown.

Here, we deploy the genetically tractable fly *D. melanogaster* to show that sublethal exposure to the pesticide emamectin benzoate (EB) suppresses female reproduction by disrupting ovulation rather than oogenesis. We identify the glutamate-gated chloride channel, GluClα, as an EB target in a neurohormonal pathway linking neural activity to octopaminergic signaling, ecdysteroid biosynthesis, and follicle rupture. These findings reveal the molecular basis of how sublethal pesticide exposure disrupts reproductive physiology by perturbing neurohormonal regulation.

## RESULTS

### Sublethal EB exposure induces strong and persistent reproductive suppression in *Drosophila*

To identify commonly used insecticides that affect female reproduction under sublethal exposure, we performed a feeding-based screening assay using a panel of compounds at 10 ppm, a concentration selected based on preliminary dose–response analyses to avoid significant mortality while allowing detection of reproductive effects (Fig. S1). Among all compounds tested, emamectin benzoate (EB) caused the most pronounced reduction in the number of eggs laid within 24 h (Figure 1A). Notably, at the tested concentration, EB did not significantly affect survival (Fig. S1), indicating that sublethal exposure is sufficient to strongly impair reproductive output.

**Figure 1.**
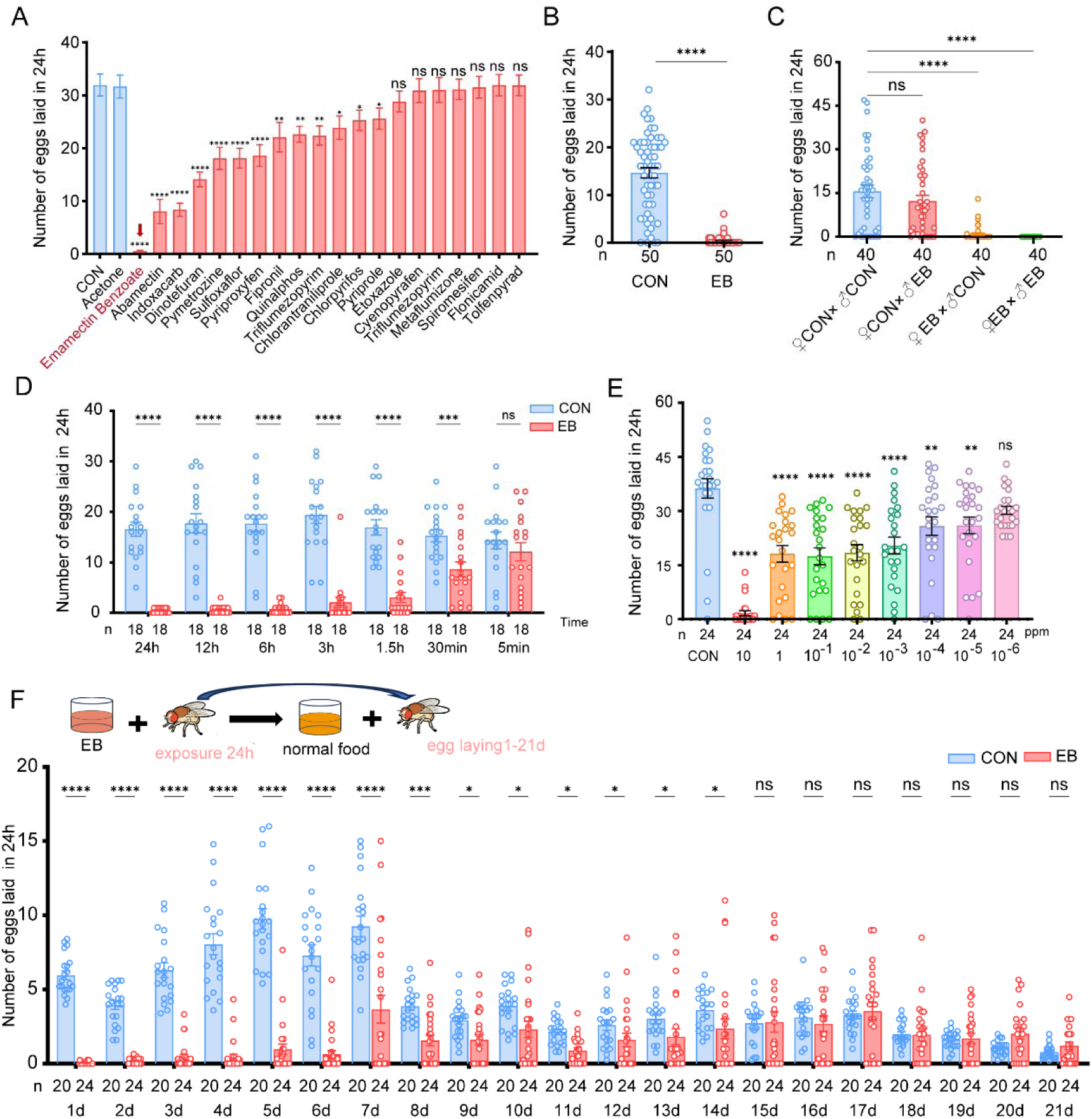
Emamectin benzoate rapidly and persistently suppresses egg laying in *Drosophila melanogaster*. (A) Number of eggs laid per female within 24 h following exposure to different insecticides. (B) Number of eggs laid per female within 24 h following exposure to sublethal emamectin benzoate (EB; LC₁, 10 ppm), as determined by bioassay. (C) Number of eggs laid per female within 24 h from different mating combinations following 10 ppm EB exposure. (D) Number of eggs laid per female within 24 h following 10 ppm EB exposure for the indicated durations. (E) Number of eggs laid per female within 24 h following exposure to EB at concentrations ranging from 10 to 10⁻L ppm. (F) Number of eggs laid per female per day after 24 h exposure to 10 ppm EB followed by transfer to normal food. Data are presented as mean ± s.e.m. Statistical significance was determined using one-way ANOVA followed by Tukey’s multiple comparison test for panels A and C–F, and Student’s *t* test for panel B. ns, not significant; *P* < 0.05 (\**), P < 0.01 (*\*\**), P < 0.001 (*\*\*\*), and *P* < 0.0001 (****)

To further define the sublethal nature of EB exposure, we determined dose–response relationships using standard bioassays (28). The concentration used in subsequent experiments (10 ppm) corresponds to a low sublethal dose, approximately LC₁ (Table. S1). We next examined the effect of EB exposure on egg laying in adult mated flies. Exposure to EB at 10 ppm significantly reduced the number of eggs laid within 24 h compared to controls (Figure 1B, Fig. S2 A). Selective exposure experiments further showed that EB treatment of females, but not males, was sufficient to reduce egg laying (Figure 1C). Consistently, EB exposure reduced egg laying in virgin females, indicating that the reproductive defects are independent of mating status (Fig. S2.C). In addition, EB exposure during larval stages did not affect adult egg laying (Fig. S2.B), suggesting that the observed phenotype arises from effects on adult physiology rather than developmental exposure.

To determine the temporal dynamics of EB-induced reproductive suppression, females were exposed to EB for varying durations. Even short-term exposure resulted in a marked reduction in egg laying, with longer exposures producing progressively stronger effects (Figure 1D). Consistently, EB reduced egg laying across a broad concentration range in a dose-dependent manner. (Figure 1E). Finally, we assessed the persistence of EB-induced reproductive defects following removal of the compound. Females exposed to EB for 24 h exhibited sustained suppression of egg laying for up to 14 days after transfer to normal food (Figure 1F), indicating that transient exposure is sufficient to induce long-lasting reproductive impairment.

### EB disrupts ovulation and egg transport without impairing oogenesis

To investigate the basis of EB-induced reproductive suppression, we first examined whether general physiological functions or ovarian development were affected by feeding flies EB. EB exposure did not alter locomotor activity, copulation rate, or early ovarian morphology in virgin females prior to egg laying (Fig. S3 A–E), indicating that neither general physiology nor early reproductive development is impaired under these conditions.

We next examined ovarian morphology in mated females following EB exposure. EB-treated females exhibited enlarged ovaries compared to controls (Figure 2A, B). This enlargement was associated with a marked accumulation of mature eggs within the ovarioles (Figure 2C, D) (25). In contrast, control ovaries contained fewer mature eggs that were more evenly distributed. Despite this accumulation, the size and morphology of individual mature eggs were similar between groups (Fig. S3.F, G), indicating that oogenesis proceeds normally under EB treatment. The increased ovarian size together with the accumulation of mature eggs suggests a defect in egg release.

**Figure 2.**
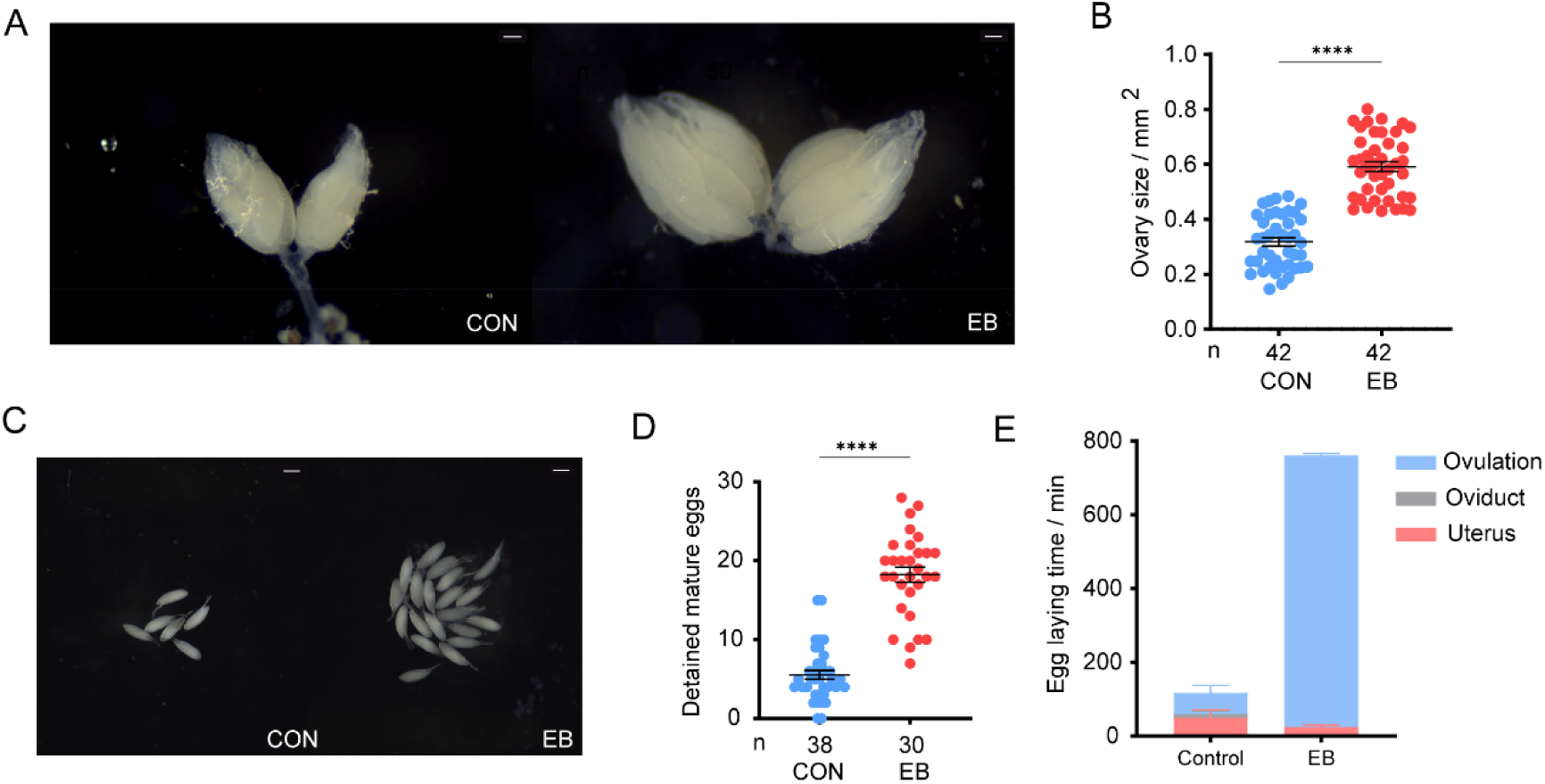
Sublethal emamectin benzoate exposure disrupts egg-laying dynamics in *Drosophila melanogaster*. (A) Representative images of ovaries from control and EB-treated females. Scale bar, 100 μm. (B) Quantification of ovary size calculated from measured length and width. (C) Representative images of isolated mature eggs from control and EB-treated females. Scale bar, 100 μm. (D) Number of detained mature eggs per female. (E) Quantification of egg distribution along the reproductive tract (ovulation, oviduct, and uterus) to estimate egg transit dynamics. Data are presented as mean ± s.e.m. Statistical significance was determined using Student’s *t*-test for comparisons between two groups (B and D). *P < 0.0001 (*\*\*\*).

To further characterize egg movement along the reproductive tract, we quantified egg distribution across the ovary, oviduct, and uterus (29). In control females, eggs were efficiently transferred from the ovary to downstream regions, whereas in EB-treated females, the majority of eggs accumulated in the ovary with minimal progression into the oviduct and uterus (Figure 2E). This altered distribution indicates a disruption in egg transit along the reproductive tract.

Together, these results indicate that EB-induced reproductive suppression primarily reflects impaired ovulation and egg transport, rather than defective oogenesis or general physiological impairment.

### GluClα-expressing neurons mediate EB-induced ovulation defects

Given that EB is known to target glutamate-gated chloride channels (GluClα) and GABA receptors (Rdl) (30, 31), we next tested whether manipulation of these molecular targets could recapitulate EB-induced reproductive phenotypes. To this end, we manipulated GluClα-expressing neurons by activating them thermogenetically with dTrpA1 or functionally silencing them by expressing the hyperpolarizing channel Kir2.1 using the GAL4-UAS system. These tools are commonly used in *Drosophila* neurogenetics to manipulate neuronal activity in defined neuronal populations (32). Both activation and functional silencing of GluClα neurons significantly reduced egg laying and increased the number of detained mature eggs, phenocopying the reproductive defects observed following EB exposure (Figure 3A–F). Dissection of ovaries revealed enlarged ovaries with accumulation of mature eggs, consistent with ovulation defects induced by EB (Figure 3A, C, D). In contrast, dTrpA1 activation of Rdl-expressing neurons did not produce comparable effects on egg laying or egg retention (Figure 3A–C), indicating that the EB-induced reproductive phenotype is specifically associated with GluClα rather than Rdl.

**Figure 3.**
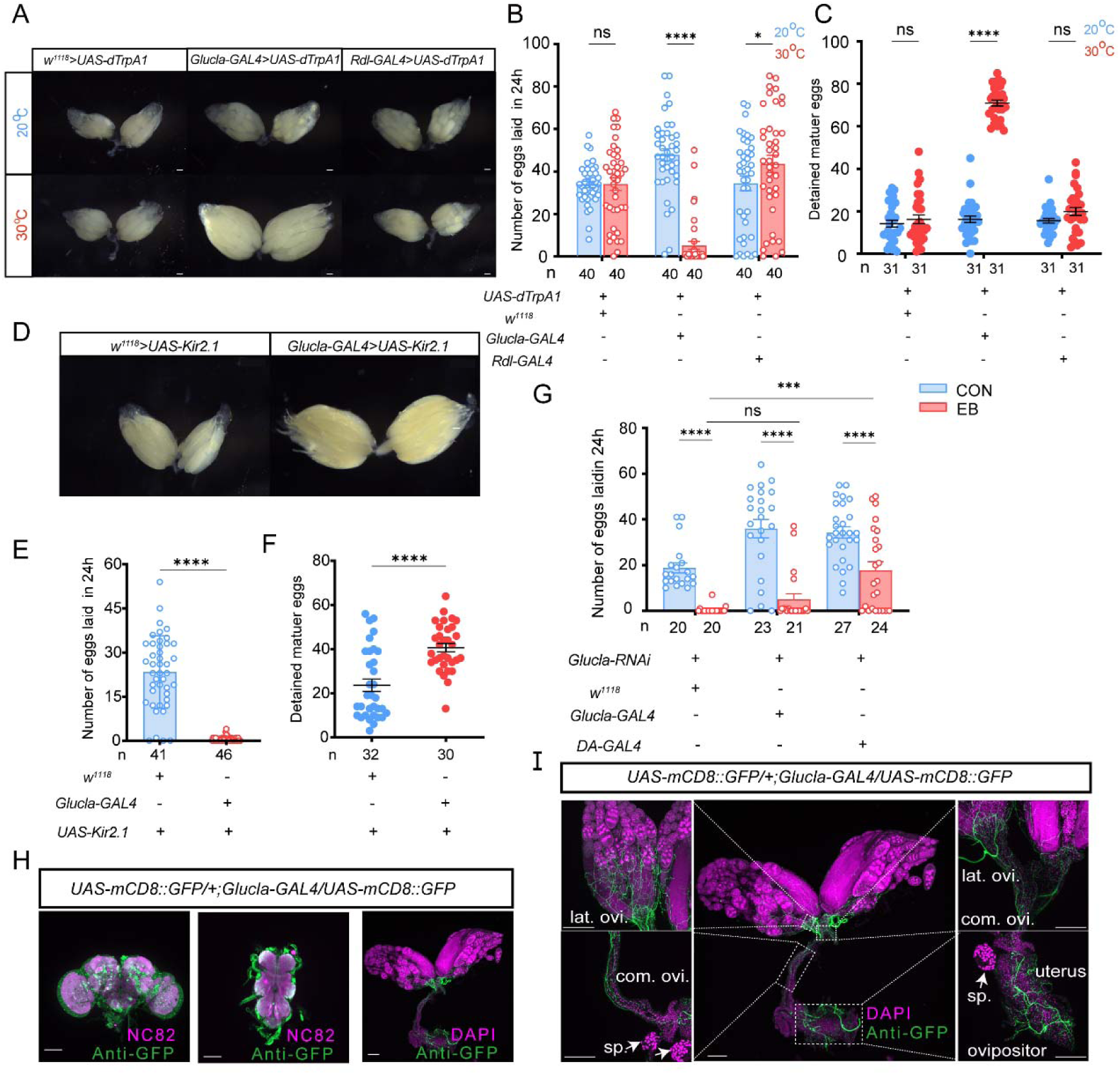
GluClα-expressing neurons mediate the egg-laying suppression induced by EB. (A) Representative ovarian morphology following thermogenetic activation (*UAS-dTrpA1*) at 20°C (control) or 30°C (activation). (B) Number of eggs laid within 24 h following neuronal activation. (C) Number of detained mature eggs following neuronal activation. (D) Representative ovarian morphology following neuronal inactivation using *UAS-Kir2.1*. (E) Number of eggs laid within 24 h following GluClα neuronal silencing. (F) Number of detained mature eggs following GluClα neuronal silencing. (G) Number of eggs laid within 24 h in control flies, GluClα knockdown flies, and flies exposed to 10 ppm EB, alone or in combination. (H) Expression pattern of *GluCl*α*-GAL4* revealed by *UAS-mCD8::GFP* labeling in the brain, ventral nerve cord (VNC), and ovary. Neural structures in the brain and VNC were labeled with anti-NC82. Ovarian nuclei were stained with DAPI. Scale bars, 100 μm. (I) Higher-magnification view of *GluCl*α*-GAL4* expression in the ovary and reproductive tract, including lateral oviduct (lat. ovi.), common oviduct (com. ovi.), uterus, spermatheca (sp.), and ovipositor. Scale bars, 100 μm. Data are presented as mean ± s.e.m. Statistical significance was determined using one-way ANOVA followed by Tukey’s multiple comparison test for multi-group comparisons (B, C, and G) and Student’s *t*-test for comparisons between two groups (E and F). ns, not significant; *P* < 0.05 (\**), P < 0.01 (*\*\**), P < 0.001 (*\*\*\*), and *P* < 0.0001 (****).

We next examined whether GluClα function contributes to EB-induced reproductive suppression. Knockdown of GluClα using the ubiquitous driver *DA-GAL4* to induce RNAi partially restored egg laying in EB-treated flies, and a similar trend toward rescue was observed using *GluCl*α*-GAL4* (33), although the effect did not reach statistical significance (Figure 3G). These results suggest that GluClα contributes to EB-induced reproductive suppression.

To determine the anatomical distribution of GluClα-expressing cells, we visualized *GluCl*α*-GAL4* expression using *UAS-mCD8::GFP*. Strong expression was observed in the brain and ventral nerve cord (VNC), where numerous neuronal cell bodies were detected. In contrast, although GFP signals were present in neuronal processes in the ovary and reproductive tract, no corresponding cell bodies were observed locally, suggesting that these processes likely represent axons derived from the central nervous system (Figure 3H; Fig. S4.A–C).

High-resolution imaging revealed that GluClα-positive axon projections extend into key regions of the reproductive tract, including the lateral oviduct, common oviduct, uterus, and ovipositor (Figure 3I). Together, these data indicate that the cell bodies of *GluCl*α-expressing neurons are located in the central nervous system and may regulate ovulation through axonal projections to the reproductive system.

### Octopaminergic GluClα neurons in the ventral nerve cord mediate ovulation suppression

To identify the specific GluClα neurons responsible for EB-induced reproductive suppression, we first used an intersectional strategy combining *Otd-nsl:FLPo* with *tub>GAL80* (34). This approach allowed us to divide GluClα neurons into brain-restricted and brain-extrinsic subpopulations, which we then manipulated separately to determine their specific contributions to EB-induced reproductive suppression. Selective activation of brain-extrinsic GluClα neurons using *UAS-dTrpA1* led to a marked reduction in egg laying and a significant accumulation of detained mature eggs, phenocopying the reproductive defects observed following EB exposure (Figure 4A–C). In contrast, activation of brain-restricted GluClα neurons did not produce significant egg-laying defects (Figure 4B, C), indicating that brain-extrinsic GluClα neurons are primarily responsible for mediating ovulation defects.

**Figure 4.**
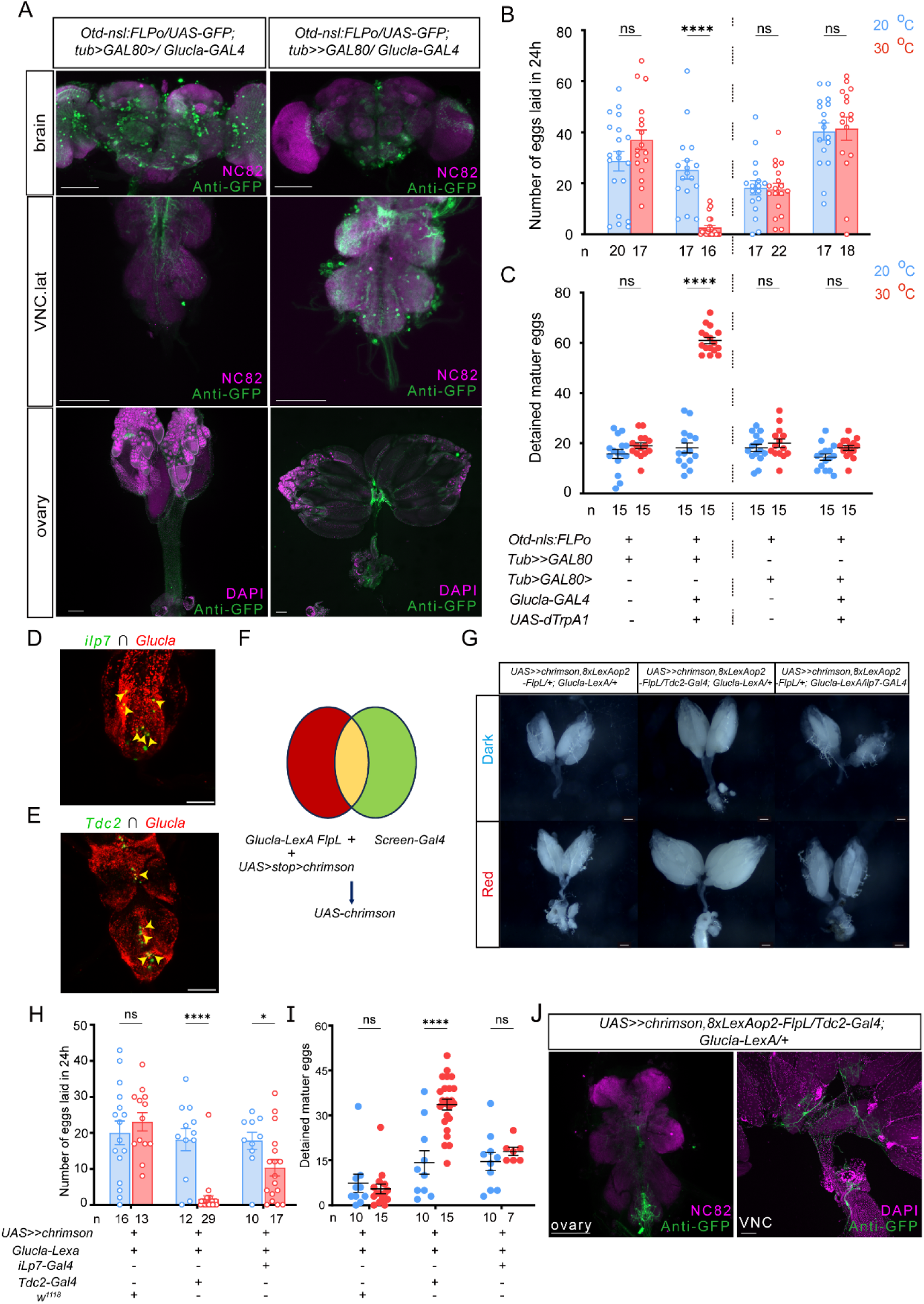
Brain-extrinsic GluClα neurons overlap with octopaminergic neurons to control egg laying. (A) Intersectional restriction of *GluCl*α*-GAL4* expression using O*td-nsl:FLPo* and *tub>GAL80*. Brain-restricted and brain-extrinsic GluClα neurons were labeled with *UAS-mCD8::GFP*. Neural structures in the brain and ventral nerve cord (VNC) were labeled with anti-NC82. Ovarian nuclei were stained with DAPI. Scale bars, 100 μm. (B) Number of eggs laid within 24 h following thermogenetic activation (20°C vs 30°C) of brain-restricted or brain-extrinsic GluClα neurons. (C) Number of detained mature eggs following activation of brain-restricted or brain-extrinsic GluClα neurons. (D–E) Co-localization of GluClα neurons with *ilp7-GAL4* (D) or *Tdc2-GAL4* (E) using intersectional labeling with *UAS-Stinger* and *LexAop-tdTomato* reporters. Genotypes: *W-; UAS-Stinger, LexAop-tdTomato/+; ilp7-GAL4/GluCl*α*-LexA*; *W-; UAS-Stinger, LexAop-tdTomato/Tdc2-GAL4; GluCl*α*-LexA/+*. Scale bars, 100 μm. (F) Schematic illustration of the intersectional activation strategy using *UAS>stop>Chrimson* and *8xLexAop2-FlpL* to selectively activate GluClα ∩ driver neurons. (G) Representative ovarian morphology following optogenetic activation of intersectional neuronal populations under dark or red-light stimulation. (H) Number of eggs laid within 24 h following optogenetic activation of intersectional neuronal populations. (I) Number of detained mature eggs following optogenetic activation. (J) Intersectional labeling of GluClα ∩ Tdc2 neurons showing projections in the ovary and reproductive tract. GFP signals were detected in the ovary and reproductive tract. Neural structures were labeled with anti-NC82 and nuclei were stained with DAPI. Scale bars, 100 μm. Data are presented as mean ± s.e.m. Statistical significance was determined using one-way ANOVA followed by Tukey’s multiple comparison test for multi-group comparisons (B, C, H, and I). ns, not significant; *P* < 0.05 (\**), P < 0.01 (*\*\**), P < 0.001 (*\*\*\*), and *P* < 0.0001 (****).

To further narrow down this population, we next tested *split-GAL4* drivers that label subsets of GluClα-expressing neurons (35, 36). However, thermogenetic activation of these split-GAL4-labeled subsets did not produce significant egg-laying defects (Fig. S5.A-B), suggesting that the critical GluClα population was not fully captured by these restricted lines.

We therefore adopted an alternative strategy to refine the candidate population by examining overlap with more restricted neuronal drivers in the ventral nerve cord. We focused on Tdc2-positive octopaminergic neurons and ilp7-positive peptidergic neurons, as these drivers label relatively small neuronal populations and have been implicated in reproductive regulation (37, 38). Co-expression analysis revealed overlap between GluClα neurons and both ilp7- and Tdc2-positive neurons (Figure 4D, E). To directly test the functional contribution of these overlapping populations, we used an intersectional optogenetic strategy to selectively activate GluClα ∩ ilp7 or GluClα ∩ Tdc2 neurons using Chrimson-mediated red-light stimulation (Figure 4F) (39). Activation of GluClα ∩ Tdc2 neurons resulted in a strong reduction in egg laying and a marked increase in detained mature eggs, closely resembling the EB-induced phenotype (Figure 4G–I). In contrast, activation of GluClα ∩ ilp7 neurons did not produce comparable effects (Figure 4H, I). Consistent with a role in peripheral reproductive control, intersectionally labeled GluClα ∩ Tdc2 neurons exhibited projections in the ovary and reproductive tract without detectable local neuronal cell bodies (Figure 4J).

Together, these results indicate that EB-induced reproductive suppression is mediated by brain-extrinsic GluClα neurons and further localize the critical functional population to a subset of octopaminergic neurons in the ventral nerve cord.

### EB impairs ovulation by suppressing OA-induced follicle rupture and ovarian ecdysteroid signaling

Given that the functionally defined GluClα ∩ Tdc2 neuronal population identified in Figure 4 is octopaminergic, and octopamine (OA) has been shown to regulate follicle rupture during ovulation (24), we next tested whether EB affects OA-dependent ovulation processes in the ovary.

We first examined ovarian morphology following EB exposure. In control females, yellow deposits corresponding to corpora lutea were readily observed at the base of the ovary, indicative of normal ovulation events. In contrast, these structures were largely absent in EB-treated females (Figure 5A), suggesting impaired ovulation and raising the possibility of defective follicle rupture (40). To directly test this, we performed ex vivo follicle rupture assays using OA stimulation, as previously described (41). OA treatment significantly promoted follicle rupture in control follicles and also increased the proportion of ruptured follicles in follicles isolated from EB-treated females. However, the level of follicle rupture in the EB + OA group remained significantly lower than that observed in OA-treated control follicles (Figure 5B, C), indicating that EB exposure compromises OA-dependent follicle rupture.

**Figure 5.**
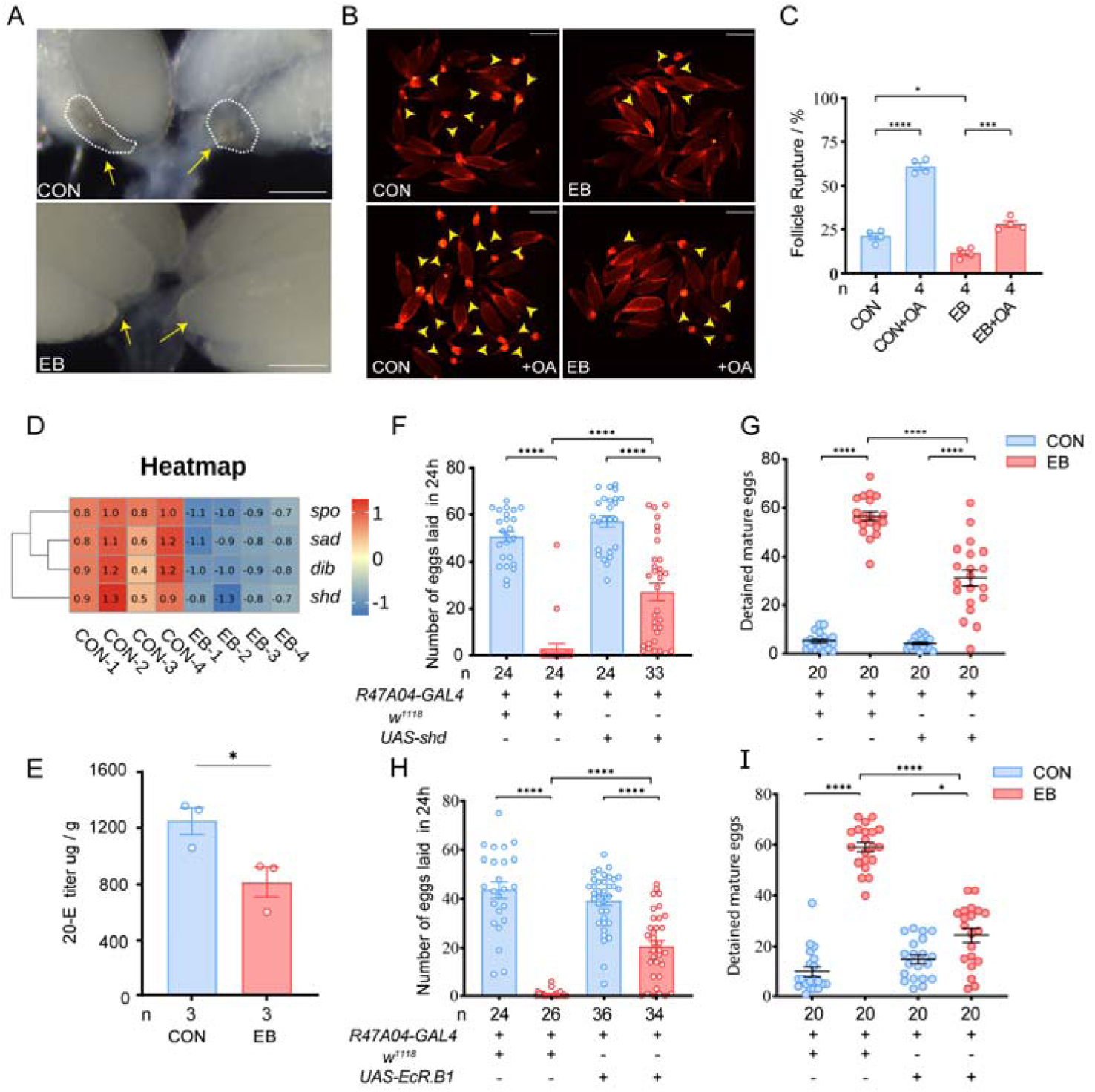
EB suppresses OA-induced follicle rupture and ovarian ecdysteroid signaling. **(A)** Representative images of dissected ovaries from control (CON) and EB-treated (EB) females. Corpora lutea deposits at the ovarian base are present in control but absent in EB-treated females. Scale bars, 100 μm. **(B)** Representative images of isolated mature follicles from control and EB-treated females cultured ex vivo in the presence or absence of octopamine (OA). Arrowheads indicate ruptured follicles. **(C)** Quantification of follicle cell rupture shown in (B). **(D)** Heatmap showing the expression levels of ecdysteroid biosynthetic and activation genes (*spo, sad, dib, shd*) in whole females following control and EB-treated females based on RNA-seq analysis. **(E)** Whole-body 20-hydroxyecdysone (20E) titers from control and EB-treated females. **(F)** Number of eggs laid within 24h in control and EB-treated flies with or without follicle cell-specific overexpression of *shd*. **(G)** Number of detained mature eggs in the same groups as in (F). **(H)** Number of eggs laid within 24 h in control and EB-treated flies with or without follicle cell-specific overexpression of *EcR.B1*. **(I)** Number of detained mature eggs in the same groups as in (H). Data are presented as mean ± s.e.m. Statistical significance was determined using Student’s *t* test for two-group comparisons (C and E) and one-way ANOVA followed by Tukey’s multiple comparison test for multi-group comparisons (F–I). ns, not significant; *P* < 0.05 (\**), P < 0.01 (*\*\**), P < 0.001 (*\*\*\*), and *P* < 0.0001 (****).

To determine whether EB acts directly on the ovary to influence follicle rupture, we first examined the tissue distribution of *GluCl*α using the FlyAtlas2 database (42). *GluCl*α expression was highly enriched in nervous tissues, including the thoracico-abdominal ganglion (TAG) and brain/CNS, whereas little or no expression was detected in reproductive tissues such as the ovary and spermatheca (Fig. S6). We next performed ex vivo assays in which isolated mature follicles were directly exposed to EB and OA. Under these conditions, EB treatment did not significantly alter OA-induced follicle rupture (Fig. S7). Together, these findings indicate that EB affects follicle rupture through upstream OA-dependent regulation.

Notably, OA stimulation did not fully restore follicle rupture in follicles isolated from EB-treated females (Figure 5B, C), raising the possibility that additional pathways contributing to ovulation are affected by EB exposure. To explore this possibility, we performed whole-body transcriptomic analysis of control and EB-treated females. Principal component analysis revealed clear separation between control and EB-treated samples (Fig. S8A). KEGG enrichment analysis identified multiple hormone-related pathways, including insect hormone biosynthesis, ovarian steroidogenesis, steroid biosynthesis, estrogen signaling pathway, and progesterone-mediated oocyte maturation (Fig. S8B). Given the enrichment of hormone-related pathways, we next examined the expression of genes involved in ecdysteroid biosynthesis and activation. Several key genes in this pathway, including *spo*, *sad*, *dib*, and *shd*, were downregulated following EB exposure (Figure 5D). Consistent with these transcriptional changes, whole-body measurement of 20-hydroxyecdysone (20E) revealed a significant reduction in ecdysteroid titers in EB-treated females (Figure 5E).

Previous studies have shown that ecdysteroid signaling in mature follicles is essential for ovulation in *Drosophila* (26). We therefore tested whether enhancing ecdysteroid signaling could alleviate EB-induced reproductive defects. Overexpression of *shade* (*shd*), which converts ecdysone into the active hormone 20E (43), partially restored egg laying in EB-treated females (Figure 5F) and reduced the accumulation of detained mature eggs (Figure 5G). Similarly, overexpression of the ecdysone receptor *EcR* in follicle cells partially rescued egg laying (Figure 5H) and ovulation defects (Figure 5I) (43).

Together, these results indicate that EB impairs ovulation through disruption of both OA-dependent follicle rupture and ovarian ecdysteroid signaling.

### EB disrupts oviduct muscle dynamics and induces conserved ovulation defects across dipteran insects

In addition to the ovulation defects described above, we next examined whether EB exposure also affects reproductive tract dynamics that may further influence egg laying efficiency.

Because successful ovulation requires coordinated muscle activity within the oviduct, we assessed contraction dynamics of the lateral oviduct following EB treatment (Figure 6A) (23, 44). Quantification revealed that EB exposure significantly increased the contraction frequency of oviduct muscles compared to controls (Figure 6B), indicating altered motor activity in the reproductive tract.

**Figure 6.**
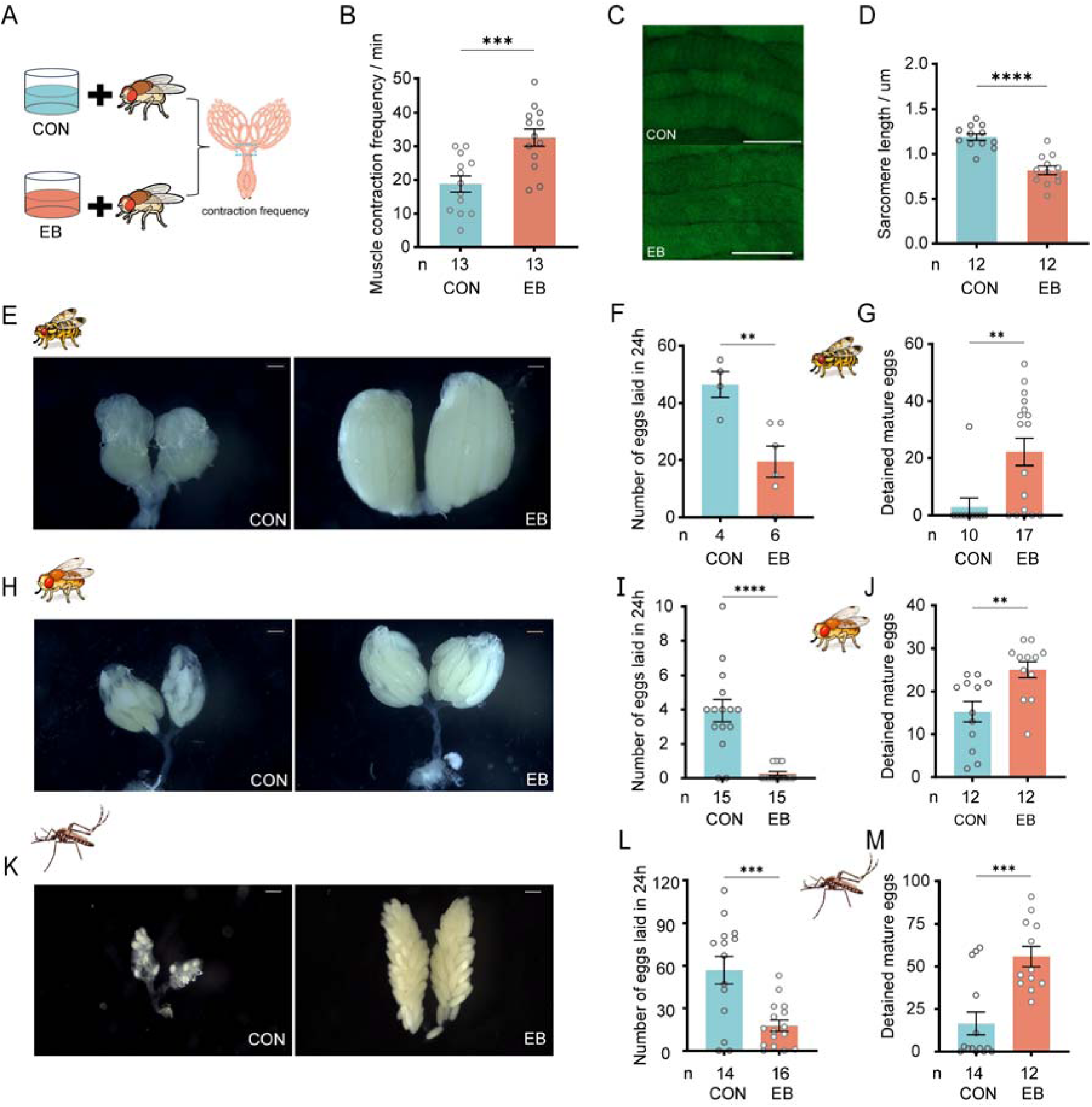
EB disrupts oviduct muscle contraction and induces conserved ovulation defects across dipteran insects. **(A)** Schematic illustration of the experimental design for measuring lateral oviduct contraction frequency in control (CON) and EB-treated females. **(B)** Quantification of oviduct muscle contraction frequency in control and EB-treated females. **(C)** Representative images of sarcomere organization in oviduct muscles labeled with MHC-GFP in CON and EB-treated flies. Scale bars, 100 μm. **(D)** Quantification of sarcomere length in oviduct muscles from control and EB-treated females. (**E**) Representative ovarian morphology in *Bactrocera cucurbitae* under control and EB treatment conditions. (**F**) Number of eggs laid within 24 h in control and EB-treated *Bactrocera cucurbitae*. (**G**) Number of detained mature eggs in control and EB-treated *Bactrocera cucurbitae*. (**H**) Representative ovarian morphology in *Drosophila suzukii* under control and EB treatment conditions. (**I**) Number of eggs laid within 24 h in control and EB-treated *Drosophila suzukii*. (**J**) Number of detained mature eggs in control and EB-treated *Drosophila suzukii*. **(K)** Representative ovarian morphology of *Aedes aegypti* under control and EB treatment conditions. **(L)** Number of eggs laid within 24 h by control and EB-treated *Aedes aegypti* females. **(M)** Number of detained mature eggs in control and EB-treated *Aedes aegypti* females. Data are presented as mean ± s.e.m. Statistical significance was determined using Student’s *t* test. ns, not significant; *P* < 0.05 (\**), P < 0.01 (*\*\**), P < 0.001 (*\*\*\*), and *P* < 0.0001 (****).

To determine whether EB directly affects lateral oviduct muscle activity, we established an ex vivo assay to measure lateral oviduct muscle contraction and used octopamine (OA) as a positive control (Fig. S9A). OA treatment robustly induced muscle contraction, whereas direct exposure to EB failed to elicit detectable contractile responses (Fig. S9B). Together, these findings indicate that EB affects oviduct muscle activity through upstream regulatory pathways.

To further evaluate muscle properties, we examined sarcomere organization in oviduct muscles using MHC-GFP labeling. In this system, the distance between GFP-labeled bands reflects sarcomere length and thus muscle tension. Consistent with previous studies showing that relaxation of oviduct muscle facilitates egg passage, EB-treated females exhibited a significant reduction in sarcomere length compared to controls (Figure 6C, D), indicating disrupted contractile dynamics of oviduct muscles (45). These results indicate that EB exposure disrupts reproductive tract muscle dynamics, which may reduce the efficiency of egg release. Together with the defects in follicle rupture described above, these findings indicate that EB disrupts multiple coordinated processes required for successful ovulation.

To determine whether EB-induced reproductive defects are conserved across insect species, we examined its effects in additional dipteran insects. In *Bactrocera cucurbitae*, EB treatment reduced egg laying and increased the number of detained mature eggs compared with controls (Figure 6E–G). Similar effects were observed in *Drosophila suzukii*, in which EB exposure led to decreased egg laying and increased egg retention (Figure 6H–J). We further tested whether this reproductive suppression extends to mosquitoes. In *Aedes aegypti*, EB-treated females also exhibited markedly reduced egg laying and increased retention of mature eggs (Figure 6K–M). These results suggest that EB-induced impairment of egg release is conserved across multiple dipteran insects, although we do not at this point know whether the mechanisms behind this EB effect are conserved.

Collectively, these findings indicate that EB primarily disrupts ovulation through coordinated effects on multiple components of the ovulation process. At the neural level, EB acts through GluClα-expressing octopaminergic neuronal circuits. At the ovarian level, EB impairs follicle rupture and reduces ecdysteroid signaling. In addition, EB alters reproductive tract muscle dynamics, which may further reduce the efficiency of egg release. Together, these results support a multi-layered mechanism underlying EB-induced reproductive suppression in *Drosophila* (Figure 7).

**Figure 7.**
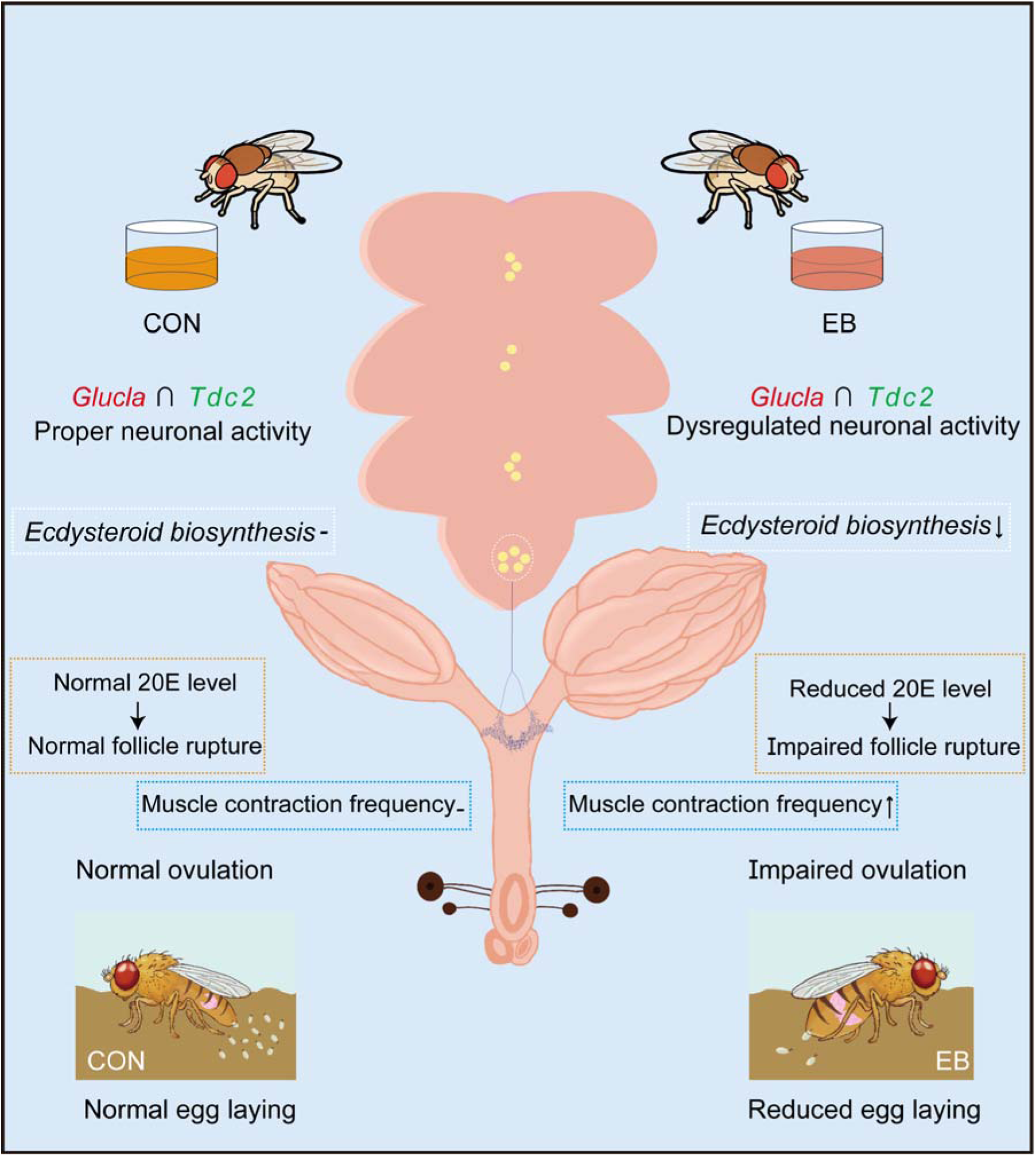
Model of EB-induced reproductive suppression in Drosophila. Schematic illustrating how sublethal emamectin benzoate (EB) exposure disrupts ovulation. Under control conditions (CON), GluClα-expressing octopaminergic (Tdc2-expressing) neurons exhibit proper activity, supporting normal ecdysteroid biosynthesis, 20-hydroxyecdysone (20E) levels, follicle rupture, and egg laying. Upon EB exposure, neuronal activity becomes dysregulated, leading to reduced ecdysteroid biosynthesis, decreased 20E levels, impaired follicle rupture, and disrupted ovulation, ultimately resulting in reduced egg laying. Arrows indicate functional relationships.

## DISCUSSION

Sublethal exposure to insecticides is increasingly recognized to have profound biological effects beyond lethality, yet the underlying mechanisms remain poorly understood (1, 3, 46). Here, we demonstrate that sublethal emamectin benzoate (EB) exposure suppresses female reproduction in *Drosophila melanogaster* by disrupting ovulation rather than oogenesis. Mechanistically, EB perturbs a specific subset of octopaminergic neurons in the ventral nerve cord, and induces defects in ecdysteroid biosynthesis, follicle rupture, and coordinated oviduct muscle contraction required for successful ovulation. These findings highlight neural regulation as a critical mediator of pesticide-induced reproductive effects and provide a framework for understanding how environmental chemicals influence fertility through neurohormonal pathways.

A key feature of this mechanism is its specificity. Rather than causing general physiological impairment, EB selectively targets GluClα-expressing neurons to disrupt reproductive output. Although both GluClα and Rdl have been proposed as molecular targets of EB, our results indicate that only GluClα-dependent pathways are functionally involved in regulating ovulation (30, 31). Furthermore, both activation and inhibition of GluClα neurons recapitulate reproductive defects, suggesting that precise regulation of neuronal activity, rather than simple gain or loss of function, is required for normal ovulation. This supports a model in which EB disrupts the balance of activity within a defined neural circuit, leading to downstream reproductive failure.

At the circuit level, we identify a subset of GluClα-expressing octopaminergic (Tdc2-expressing) neurons as a key functional population. Octopamine is well established as a central regulator of ovulation in insects (21–24), particularly through its role in promoting follicle rupture (41). Our findings extend this framework by implicating GluClα-expressing octopaminergic neurons in pesticide-induced reproductive suppression, thereby linking an ion channel target of insecticides to a defined neuromodulatory population involved in reproductive physiology. In addition to defects in follicle rupture, EB exposure also alters oviduct muscle dynamics, suggesting that successful ovulation requires coordinated regulation across neural, endocrine, and muscular levels.

In contrast to our previous findings that EB stimulates reproduction in brown planthoppers via GluClα-dependent juvenile hormone (JH) signaling, we show here that EB suppresses reproduction in *Drosophila melanogaster* through pathways involving GluClα and 20E signaling (47, 48). This contrast highlights species-specific differences in reproductive responses to EB and suggests that a common molecular target can engage distinct neurohormonal outputs in different insects. Such differences likely reflect divergence in how GluClα-dependent neural activity interfaces with endocrine regulation, ultimately leading to opposite reproductive outcomes. These findings provide a conceptual framework for understanding how insecticide-induced reproductive effects vary across species through differential organization of neurohormonal signaling.

Importantly, our findings are not restricted to *Drosophila melanogaster*. Similar reproductive suppression has been observed in multiple insect species, including tephritid flies and mosquitoes, suggesting that the inhibitory effects of EB on reproductive physiology are conserved in at least a subset of insects (49, 50). This conservation highlights ovulation and its underlying neurohormonal regulation as a vulnerable target of chemical perturbation. From an applied perspective, such effects may be exploited for pest management, as sublethal interference with reproductive processes could reduce population growth without requiring acute toxicity. These results therefore provide a mechanistic basis for developing more targeted and sustainable strategies for insect control (51).

### Limitations and future directions

Several mechanistic questions remain to be addressed. Although our results implicate GluClα-expressing octopaminergic neurons in EB-induced ovulation defects, how these neurons communicate with the ovary to regulate follicle rupture remains unclear. In particular, the ovarian receptors, target cell types, and downstream signaling events that connect octopaminergic input to follicle cell rupture require further investigation. It also remains unresolved whether GluClα-positive octopaminergic neurons directly modulate ovarian ecdysteroid-producing cells or indirectly affect ecdysteroid biosynthesis through systemic endocrine pathways or reproductive tract activity. In addition, the cellular origin and temporal dynamics of ovarian ecdysteroid synthesis during EB exposure remain to be defined. Our transcriptomic analyses also identified additional differentially expressed genes that may contribute to EB-induced reproductive defects, and their functional roles should be validated in future studies. Future work integrating neuronal activity imaging, cell-type-specific genetic manipulation, ovarian single-cell transcriptomics, and spatial profiling of ecdysteroid signaling will help clarify how neural, endocrine, and muscular outputs are coordinated during ovulation.

Together, our findings suggest that sublethal EB exposure does not simply induce general toxicity, but instead disrupts the neurohormonal networks that coordinate reproduction. By engaging GluClα-associated neural circuits and downstream endocrine and muscular outputs, EB perturbs reproductive control at multiple levels. These findings reveal ovulation as a sensitive physiological process affected by sublethal insecticide exposure and provide a foundation for understanding how environmental chemicals reshape fertility through neurohormonal mechanisms.

## Materials and Methods

### *Drosophila* Strains

*Drosophila melanogaster* strains used in this study included *Canton-S* (wild-type), *w^1118^*, *UAS-dTrpA1*, *UAS-Kir2.1*, *UAS-mCD8::GFP*, and *UAS-RedStinger*, which were maintained and expanded in our laboratory. *GluCl*α*-GAL4* and *GluCl*α*-LexA* lines were obtained from Dr. Yi Rao (Peking University, China). *MHC-GFP* and *R47A04-GAL4* lines were provided by Dr. Jian-hua Huang (Zhejiang University, China). *Otd-nsl:FLPo; tub>GAL80*, *Otd-nsl:FLPo; tub>>GAL80*, *w^−^; 8xLexAop2-FlpL*, *UAS>>Chrimson*, *UAS-RFP*, and *Ilp7-GAL4* were obtained from Dr. Yu-feng Pan (Southeast University, China). *Split-GAL4* lines *GluCl*α*-SS1* (BL86650), *GluCl*α*-SS2* (BL88524), and *GluCl*α*-SS3* (BL88335), as well as *UAS-shd* (BL60639) and *UAS-EcR.B1* (BL6969), were obtained from the Bloomington Drosophila Stock Center. All flies were reared at 25°C, 60% relative humidity, under a 12 h light/12 h dark cycle on standard *Drosophila* medium containing sugar and yeast.

### Bioassays

Adult bioassays of *Drosophila* were performed as previously described (52). Stock solutions of emamectin benzoate (EB; 95.2% active ingredient; Hebei Weiyuan Co., Ltd., China) were prepared in acetone and diluted to the desired concentrations. For bioassays, 1.5% agar containing 1% sucrose was poured into standard Drosophila vials and allowed to solidify.

Subsequently, 100 μL of EB solution was evenly applied to the agar surface, and the vials were covered with gauze to allow solvent evaporation at room temperature. Three- to five-day-old adult flies, previously starved for 4 h, were transferred to EB-treated vials at a density of 10 flies per vial, with four biological replicates for each concentration. All vials were maintained at 25°C, 60% relative humidity, under a 12 h light/12 h dark cycle. After 24 h, mortality was recorded; flies that were unable to walk normally were scored as dead. Survival data were analyzed using POLO Plus software (52).

For larval bioassays, we adopted a method described previously in our lab with minor modifications (53). Briefly, forty first instar larvae were transferred to vials containing standard fly food supplemented with EB at different concentrations. Larval mortality was recorded after 24 h under the same conditions as described above.

Sublethal concentrations (e.g., LC₁ and LC_30_) were calculated based on the dose–response data using POLO Plus, and the corresponding sublethal concentrations were used for subsequent fecundity assays.

### Fecundity Measurement

For fecundity assays, EB-treated food was prepared based on the bioassay results and allowed to dry completely. Three- to five-day-old adult females, fully mated and starved for 4 h prior to the assay, were exposed to EB-containing food for 24 h. Each treated female was then transferred individually to an oviposition chamber, and the number of eggs laid over 24 h was recorded (54).

For larval exposure experiments, females exposed to EB during the larval stage were collected after adult eclosion and paired with untreated males to assess fecundity. For assays measuring unproductive egg laying, 14-day-old virgin females were used. For experiments involving different mating combinations, 3-day-old unmated flies were selected and paired as indicated. All other procedures were performed as described above. For experiments involving different exposure durations or EB concentrations, all procedures were identical except for the specified treatment time or concentration.

Fecundity was assessed at the level of individual animals, with each data point representing the number of eggs laid by a single female. Unless otherwise specified, only adult females were treated and used for fecundity measurements.

### Ovary Size and Mature Egg Measurements

Ovaries were collected from EB-treated and control females following fecundity assays.

Images were acquired using a Motic imaging system equipped with a digital camera and Motic Images Plus 3.0 software. The number of mature eggs within each female was counted. Ovary area, as well as the length and width of mature eggs, were measured using FIJI software.

### Mating and Climbing Assays

For mating assays, adult female flies pre-treated with EB were briefly anesthetized on ice and individually transferred into a custom-designed mating chamber. Males and females were separated into two layers using a removable transparent barrier to prevent premature contact. After all flies were loaded, the chambers were placed in an incubator at 25°C, 60% relative humidity, under standard light conditions and allowed to acclimate for approximately 1 h. The barrier was then removed to allow interaction between males and females. Mating behavior was recorded using an FDR-AX30 video camera, and the percentage of successful copulation events within 1 h was quantified (55).

For climbing assays, flies were divided into three groups: EB-treated, acetone-treated (solvent control), and untreated controls. After treatment, groups of 10 flies were transferred into vertical tubes (19 cm in height, 2 cm in diameter), with 10 biological replicates per condition. Flies were gently tapped to the bottom of the tube, and climbing activity was recorded for 30 s using an FDR-AX30 video camera. The number of flies reaching the upper half of the tube was recorded at 2 s intervals. Climbing performance was quantified as the percentage of flies reaching the upper half of the tube (56).

### Egg-Laying Time Assay

Egg laying time was measured following previously reported methods with minor modifications (29). After 24 hours of EB treatment, the treated female flies were transferred to fresh standard food for 6 hours. The flies were then placed in a −80°C freezer for 5 minutes to preserve any eggs present in the reproductive tract. The location of the eggs within the reproductive tract was recorded. The egg laying time was categorized into three phases: ovulation time, oviduct time, and uterus time. Ovulation time refers to the period from egg release from the ovaries to its entry into the oviduct. Oviduct time is the duration the egg spends in the oviduct, while uterus time refers to the time spent in the uterus. The distribution of eggs in these phases was assessed by calculating the percentage of females with eggs present in either the oviduct or uterus. The egg laying time was calculated using the following formula:

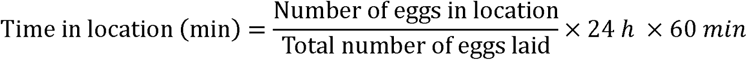

### dTrpA1 Activation and Kir2.1 Neuronal Silencing

dTrpA1 activation experiments were performed using flies carrying UAS-dTrpA1, which were maintained at 20°C to prevent premature channel activation. Female flies aged 3–5 days were pre-fed with wet yeast prior to the assay. Flies were briefly anesthetized on CO_2_ and transferred to oviposition chambers, where they were allowed to recover at 20°C for at least 30 min. The chambers were then shifted to 30°C to activate dTrpA1, while control groups were maintained at 22°C. Egg laying was quantified after 24 h.

Kir2.1 Neuronal silencing experiments were conducted using flies expressing UAS-Kir2.1, which were reared under standard conditions at 25°C. Female flies aged 3–5 days were pre-fed with wet yeast and subjected to oviposition assays. Egg laying was quantified after 24 h.

### RNAi-Mediated Knockdown and Overexpression Experiments

Gene knockdown and overexpression rescue experiments were carried out using the GAL4/UAS system. Female flies aged 3–5 days were collected after eclosion and pre-fed with wet yeast prior to the assay. Flies were exposed to EB for 24 h as described above and then individually transferred to oviposition chambers for a further 24 h. Egg laying was recorded at the end of the assay. Appropriate genetic controls were included in parallel.

### Optogenetic Stimulation

For optogenetic stimulation experiments, newly eclosed flies (1–3 days old) were collected and transferred to standard food supplemented with 0.2 mM all-trans-retinal (116-31-4, Sigma-Aldrich). Vials were wrapped in aluminum foil to maintain dark conditions, and flies were reared for 3–4 days prior to the assay.

Flies were then transferred to oviposition chambers as described above. Red light stimulation (620 nm, 0.03 mW/mm²; Shanghai Wanqi Technology Co., Ltd.) was applied continuously throughout the assay period. Egg laying was quantified after 24 h. Following the assay, ovaries were dissected for mature egg quantification. Light intensity was measured using a power meter (PS-310 V2, Gentec, Canada).

### Immunohistochemistry and Confocal Imaging

Brains, ventral nerve cords (VNC), and ovaries were dissected from 4–8-day-old adult females in 1×PBS. Tissues were fixed in 4% PFA at room temperature for approximately 25 min, followed by four washes in PAT3 (PBS containing 0.5% Triton X-100 and 0.5% bovine serum albumin), 10 min each. Samples were then incubated in blocking buffer (ReadyProbes™ Normal Goat Serum, 2.5%) at room temperature for 60 min. Primary antibodies were applied at room temperature for 4 h and then incubated overnight at 4°C. After four 10-min washes in PAT3 buffer, tissues were incubated with secondary antibodies at room temperature for 4 h, followed by four additional 10-min washes. For nuclear staining, samples were incubated with DAPI (C0060, Solarbio, 1:1000) at room temperature for 15 min. Tissues were mounted using antifade mounting medium (Vector Laboratories).

Primary antibodies and dilutions were as follows: nc82 (1:1000, Developmental Studies Hybridoma Bank, mouse), GFP (A11120, Thermo Fisher Scientific, mouse), GFP (A11122, Thermo Fisher Scientific, rabbit), and RFP (600-401-379, Rockland, rabbit, 1:1000). Secondary antibodies included Alexa 488-conjugated donkey anti-rabbit (R37118, Thermo Fisher Scientific, 1:500), Alexa 488-conjugated donkey anti-mouse (R37114, Thermo Fisher Scientific, 1:500), Alexa 555-conjugated donkey anti-rabbit (R37119, Thermo Fisher Scientific, 1:500), and Alexa 555-conjugated donkey anti-mouse (R37115, Thermo Fisher Scientific, 1:500).

Confocal imaging was performed using a Zeiss LSM980 microscope (Jena, Germany) at 10×, 20×, or 40× magnification with a resolution of 1024 × 1024 pixels or higher. Image processing and analysis were conducted using FIJI software. For TdTomato and RedStinger experiments, anti-GFP staining was omitted, and imaging was performed directly on fixed and washed samples.

### Follicle Rupture Assay

Follicle rupture assays were performed based on previously described methods with minor modifications(57). Depending on the experimental design, mature follicles were isolated either from control and EB-treated females or directly exposed to EB during ex vivo culture. Ovaries were rapidly dissected in Grace’s insect medium (1160594, Thermo Fisher Scientific) using a 9-well dissection dish (71563-01, Corning). Mature follicles were manually isolated from the ovaries and identified based on the fluorescent reporter signal (*R47A04-GAL4 > UAS-RFP*). For each replicate, 20–30 mature follicles were collected, with five biological replicates per condition.

Follicles were then incubated in Grace’s medium containing 20 μM octopamine (68631, Sigma-Aldrich) in the dark for 4 h. Following incubation, the follicle rupture rate was quantified as the percentage of ruptured follicles relative to the total number of follicles. Rupture was defined by the release of the oocyte from the follicle and visible disruption of the follicle wall.

### RNA Sequencing and Data Analysis

Total RNA was extracted using FreeZol Reagent (R711-01, Nanjing Vazyme Biotech Co., Ltd.) according to the manufacturer’s instructions. RNA samples were submitted to Guangzhou Aozhi Biotechnology Co., Ltd.. for library construction and high-throughput sequencing. Raw sequencing reads were subjected to quality control using *fastp* to remove low-quality reads and generate clean reads. Clean reads were first aligned to the Drosophila reference genome NCBI:GCF_000001215.4 using HISAT2 to assess mapping efficiency and genomic distribution. Subsequently, clean reads were aligned to the reference transcriptome using Bowtie2, and gene and transcript expression levels were quantified using RSEM. Gene expression levels were represented as both raw read counts and fragments per kilobase of transcript per million mapped reads (FPKM). These data were used to assess gene expression profiles across different samples.

### Determination of 20-Hydroxyecdysone (20E) Content by HPLC–MS/MS

Female flies were treated with EB as described above, followed by a 24 h oviposition period prior to sample collection. 20E levels were measured using HPLC–MS/MS. For each condition, three biological replicates were prepared, with 50 flies per replicate. Samples were weighed and homogenized in 400 μL methanol using a tissue grinder (65 Hz, 180 s), followed by ultrasonic extraction at 4°C for 30 min. Subsequently, 400 μL n-hexane was added, and samples were vortexed thoroughly. After centrifugation at 12,000 rpm for 15 min at 4°C, the supernatant was collected and evaporated to dryness using a vacuum concentrator. The residue was reconstituted in 200 μL methanol, vortexed, and centrifuged again at 12,000 rpm for 15 min at 4°C. The resulting supernatant was subjected to LC–MS/MS analysis. Chromatographic separation was performed on an Acquity UPLC BEH C18 column (1.7 μm, 2.1 × 100 mm) using a Waters Acquity UPLC system coupled to an AB SCIEX 5500 triple quadrupole mass spectrometer. The column temperature was maintained at 40°C, and the flow rate was set to 0.300 mL/min. A six-point standard curve was generated using 20E standards (Minxin Biotechnology Co., Ltd., Shanghai, China) for quantification.

### Muscle Contraction Assay

Muscle contraction assays were performed following EB treatment as described above. Female flies were dissected in Grace’s insect medium, and oviduct muscle contractions were recorded after a 30 s stabilization period. Videos were recorded for 1 min, and contraction frequency was manually quantified.

For ex vivo assays, ovaries were rapidly dissected in HL3.1 buffer and transferred to 90 μL of fresh HL3.1 buffer in a clean dish. Videos were recorded for 10 min to establish baseline contractions. Subsequently, 10 μL of 10 mM OA, EB, or vehicle control was slowly added to the dish to avoid mechanical disturbance, and muscle contractions were recorded for an additional 5 min. Contraction frequency was manually quantified for the post-treatment period. Each experiment included at least three independent biological replicates.

### Sarcomere Length Measurement

Sarcomere length of oviduct muscles was measured using MHC-GFP flies. Female reproductive tracts from EB-treated and control flies were dissected and immediately fixed in 4% paraformaldehyde at room temperature for 25 min. Samples were then washed four times in 1× PBS for 10 min each. Oviduct muscles labeled with GFP were imaged using confocal microscopy. Sarcomere length was determined by measuring the distance between adjacent GFP-positive bands. For each reproductive tract, three myofibrils were analyzed, and at least three sarcomeres per myofibril were measured and averaged. Four biological replicates were included for each condition.

### Reproductive assays in other insect species

For *Drosophila suzukii*, bioassays and oviposition assays were performed using the same procedures described for *Drosophila melanogaster*.

For *Bactrocera cucurbitae*, fully mated adult females were exposed to EB using the glass vial residual film method. Briefly, glass vials were coated with the indicated concentration of EB and allowed to dry completely before use. Females were exposed for 24 h and then transferred to oviposition chambers. Oviposition assays were conducted using groups of five females, and the total number of eggs laid per group was recorded after 24 h. Following egg collection, females were dissected and the number of retained mature eggs was quantified individually.

For *Aedes aegypti*, fully blood-fed females were provided with sucrose solution containing EB for 72 h. Individual females were then transferred to Petri dishes containing moist filter paper as an oviposition substrate. Eggs were counted after 24 h, and ovaries were subsequently dissected to quantify retained mature eggs. Egg laying and retained mature eggs were analyzed on an individual-female basis.

## Acknowledgments

This research was funded by the National Natural Science Foundation of China (32022011 & 32472542), the National Key Research and Developmental Program of China (2025YFE0210000) and the Fundamental Research Funds for Central Universities of the Central University (KJJQ2026014). For the purpose of open access, the author has applied a ‘Creative Commons Attribution (CC BY) licence to any Author Accepted Manuscript version arising from this submission.

## Supporting Information

### Sublethal emamectin benzoate exposure disrupts ovulation through a neurohormonal pathway in *Drosophila* and mosquitoes

**Fig. S1.**
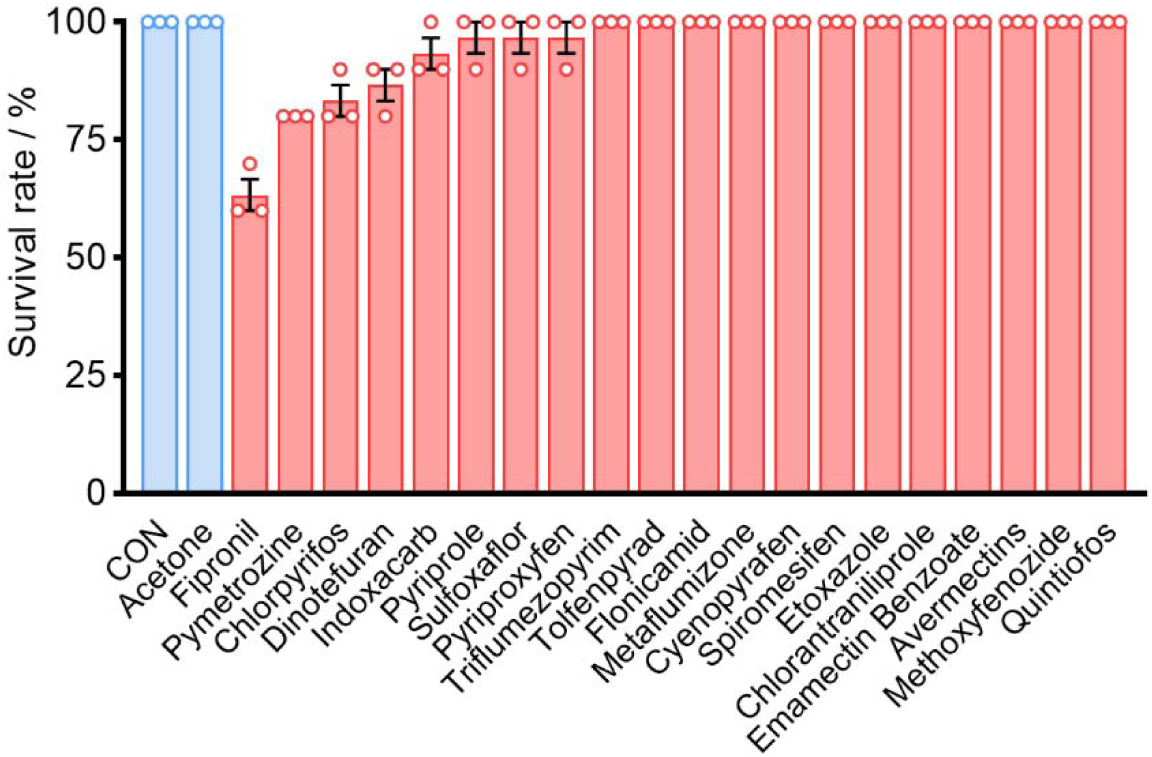
Adult female flies were exposed to 10 ppm of 17 commonly used insecticides, and survival rate was evaluated after treatment. Data represent mean ± s.e.m. from three independent biological replicates (n = 3 per replicate).

**Fig. S2.**
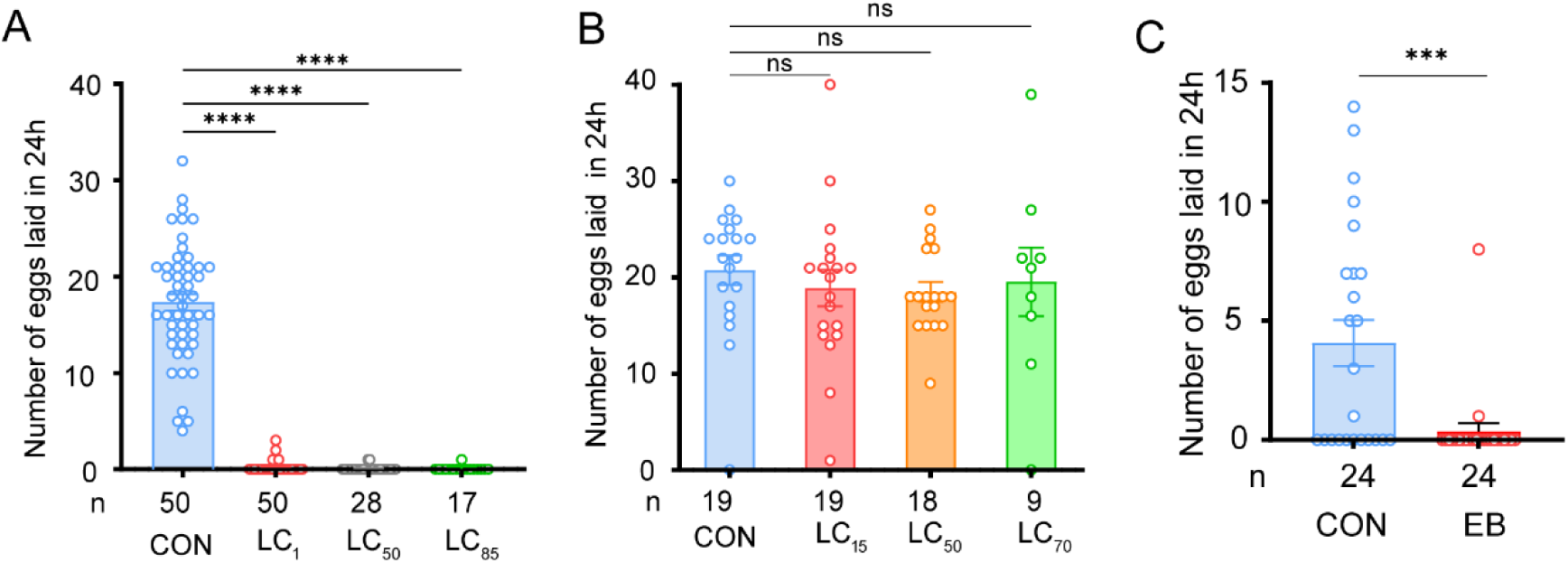
Dose-dependent effects of emamectin benzoate exposure on adult egg production and developmental stage specificity. (A) Number of eggs laid per female within 24 h following adult exposure to increasing concentrations of EB; LC₁, LC₅₀, and LC₈₅, as determined by bioassay. (B) Number of eggs laid per female within 24 h in adults that had been exposed to EB during the larval stage at LC₁₅, LC₅₀, or LC₇₀ concentrations. (C) Number of eggs laid per female within 24 h following exposure to EB in 14-day-old virgin females. Data are presented as mean ± s.e.m. Statistical significance was determined using one-way ANOVA followed by Tukey’s multiple comparison test for panels A and B, and Student’s *t* test for panel C. ns, not significant; *P* < 0.05 (\**), P < 0.01 (*\*\**), P < 0.001 (*\*\*\*), and *P* < 0.0001 (****).

**Fig. S3.**
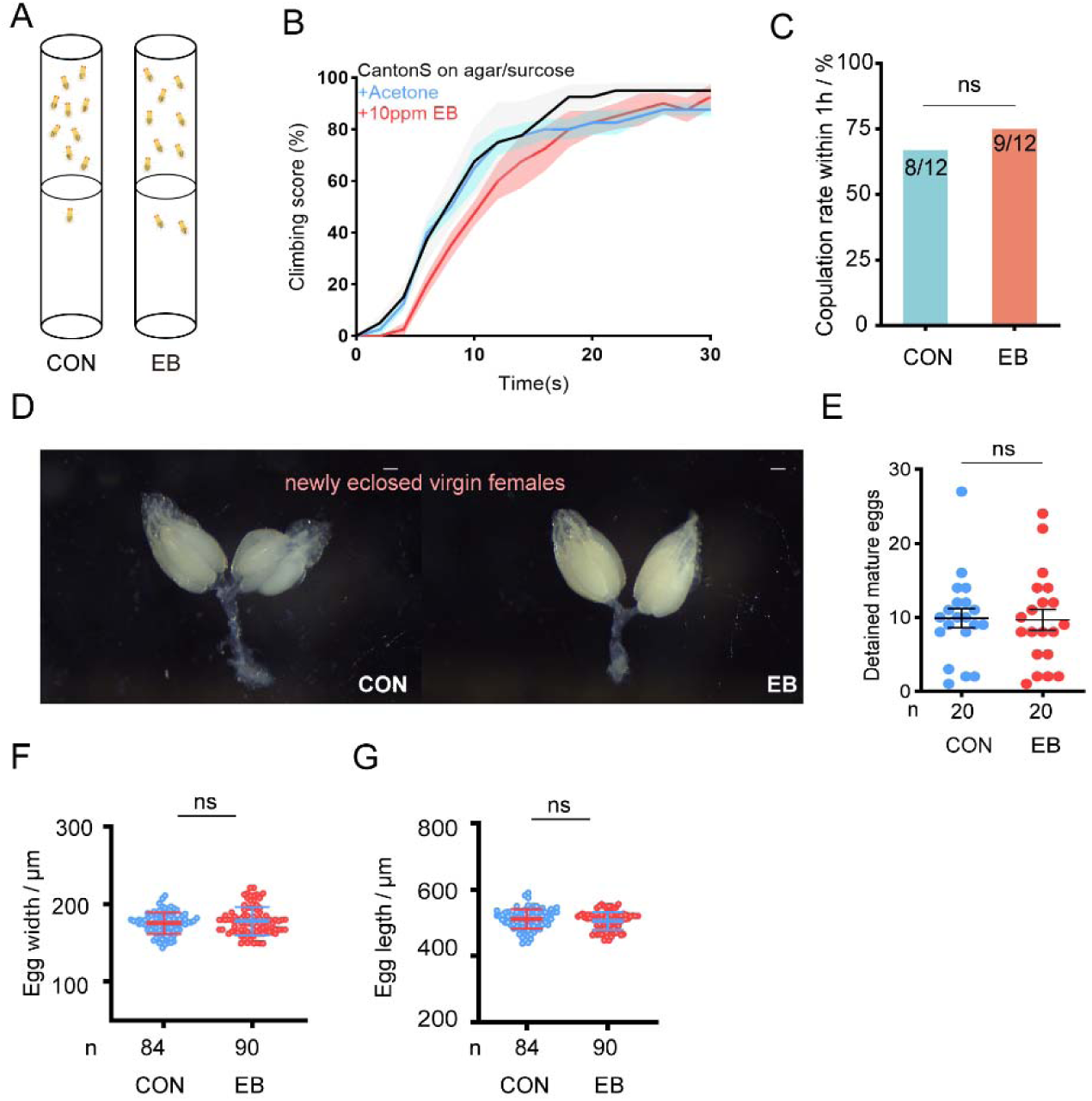
Sublethal EB exposure does not impair locomotion, mating, or ovarian development. (A) Schematic illustration of the negative geotaxis (climbing) assay. (B) Climbing performance of control flies, acetone-treated flies, and flies exposed to 10 ppm EB. Climbing score was recorded over 30 s. (C) Copulation rate within 1 h following 10 ppm exposure in virgin females (CON, 8/12; EB, 9/12). (D) Representative images of ovaries from 3–5-day-old virgin females following 10 ppm exposure. Females had not initiated egg laying at the time of dissection. Scale bar, 100 μm. (E) Number of mature eggs detained in virgin females. (F) Egg width measurements of detained mature eggs. (G) Egg length measurements of detained mature eggs. Data are presented as mean ± s.e.m. Statistical significance was determined using Student’s *t*-test for comparisons between two groups (C, E, F, and G). ns, not significant.

**Fig. S4.**
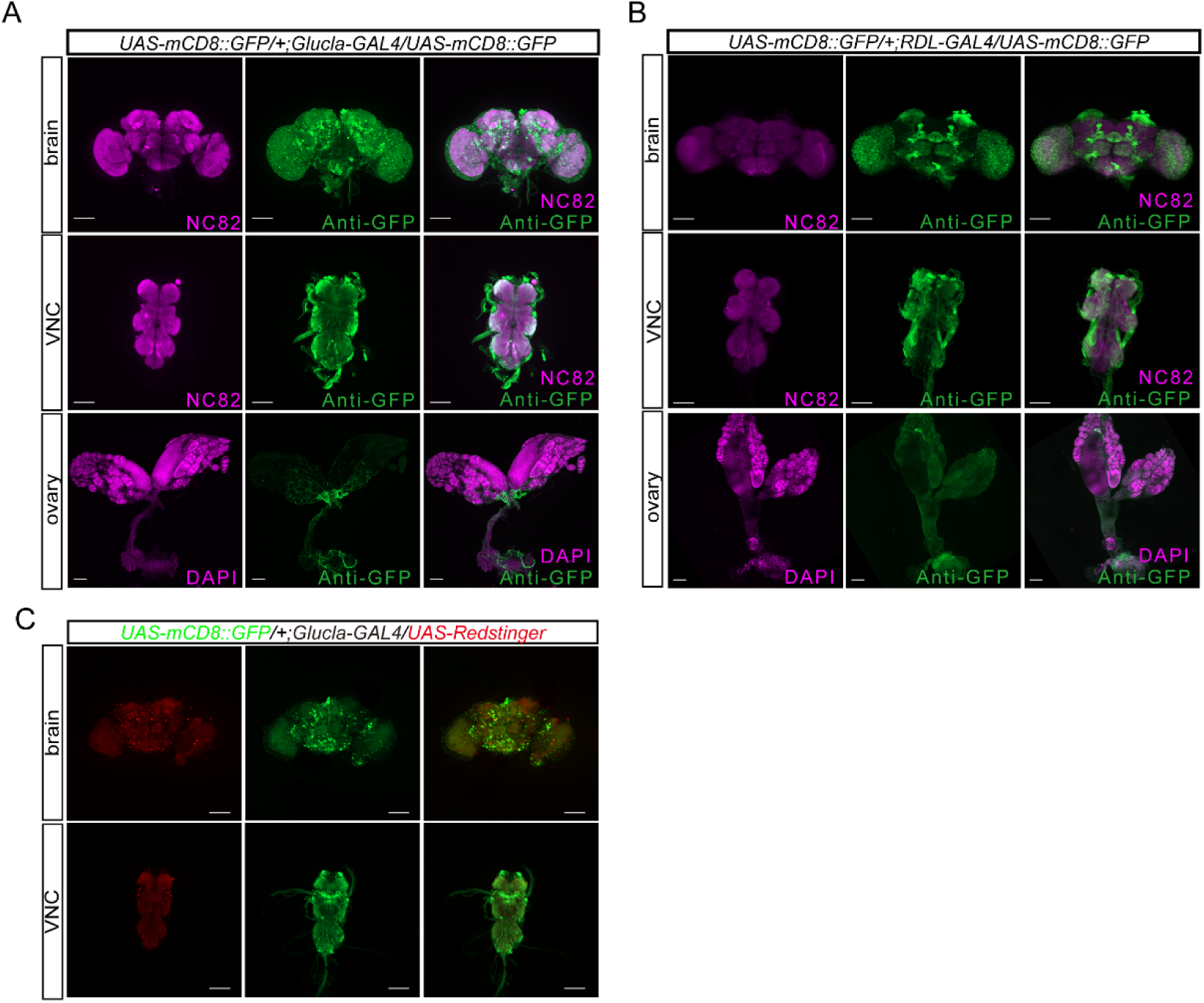
Expression patterns of *GluCl*α*-GAL4* and *Rdl-GAL4* in the nervous system and ovary. (A) Expression pattern of *GluCl*α*-GAL4* revealed by *UAS-mCD8::GFP* labeling in the brain, ventral nerve cord (VNC), and ovary. Neural structures in the brain and VNC were labeled with anti-NC82. Ovarian nuclei were stained with DAPI. GFP signals were detected using anti-GFP immunostaining. (B) Expression pattern of *Rdl-GAL4* revealed by *UAS-mCD8::GFP* labeling in the brain, VNC, and ovary. Neural structures in the brain and VNC were labeled with anti-NC82. Ovarian nuclei were stained with DAPI. (C) Co-labeling with *UAS-RedStinger* to visualize nuclei in *GluCl*α*-GAL4*–expressing cells. RedStinger-positive nuclei were observed in the brain and VNC but not within ovarian tissue. Scale bars, 100 μm.

**Fig. S5.**
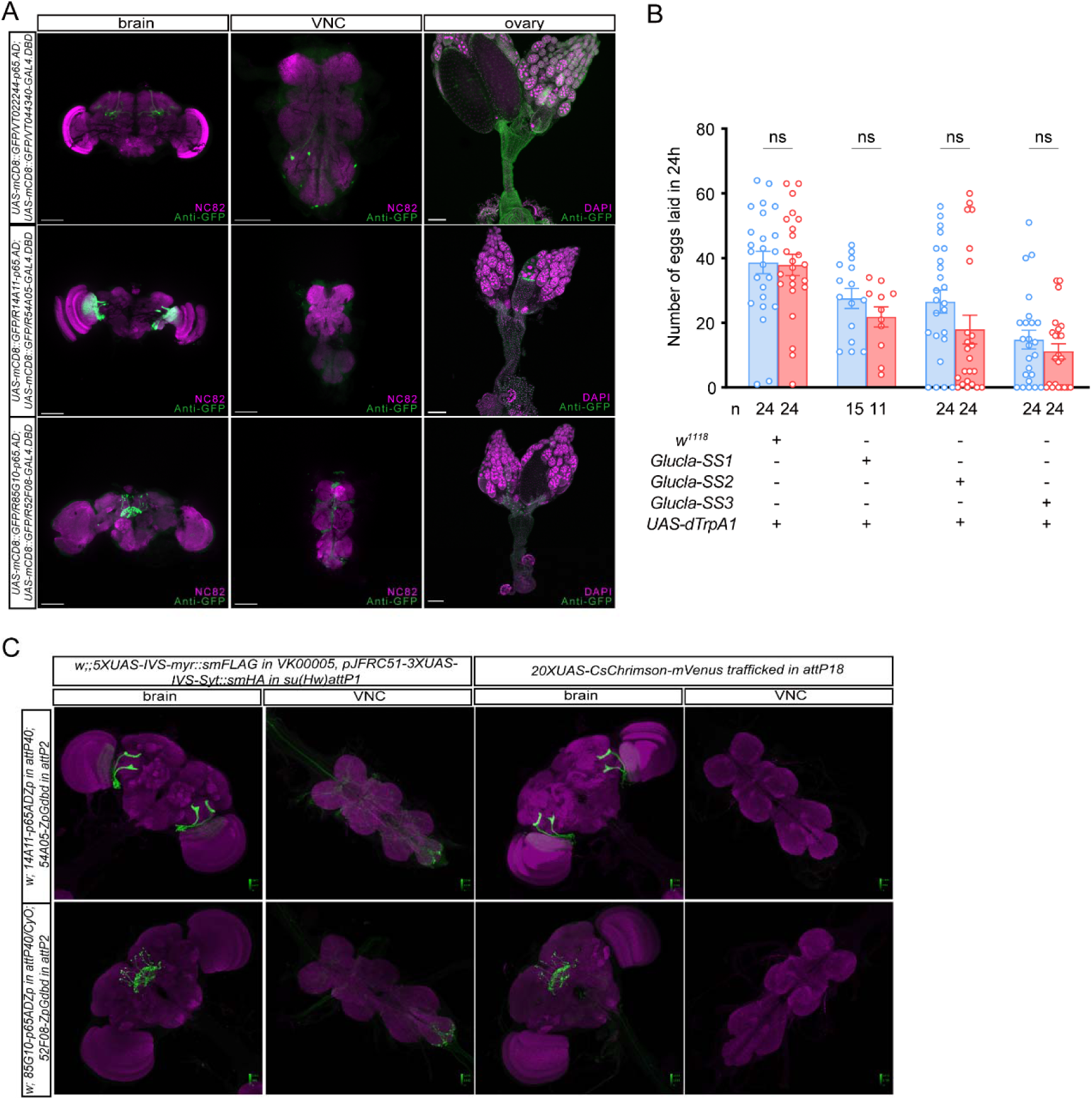
Expression patterns and functional testing of split-GAL4 drivers targeting subsets of GluClα neurons. (A) Expression patterns of three split-GAL4 drivers targeting subsets of GluClα neurons (hereafter referred to as GluClα-SS1, GluClα-SS2, and GluClα-SS3) visualized using UAS-mCD8::GFP. Neural structures in the brain and ventral nerve cord (VNC) were labeled with anti-NC82. Ovarian nuclei were stained with DAPI. Scale bars, 100 μm. (B) Number of eggs laid within 24 h following thermogenetic activation (UAS-dTrpA1; 20°C vs 30°C) of GluClα-SS1, GluClα-SS2, and GluClα-SS3. (C) Expression patterns of the corresponding split-GAL4 lines obtained from the Janelia FlyLight database using alternative reporter constructs. Images are reproduced from publicly available FlyLight resources and are shown for comparison with panel (A). Data are presented as mean ± s.e.m. Statistical significance in (B) was determined using two-way ANOVA followed by Tukey’s multiple comparison test. ns, not significant.

**Fig. S6.**
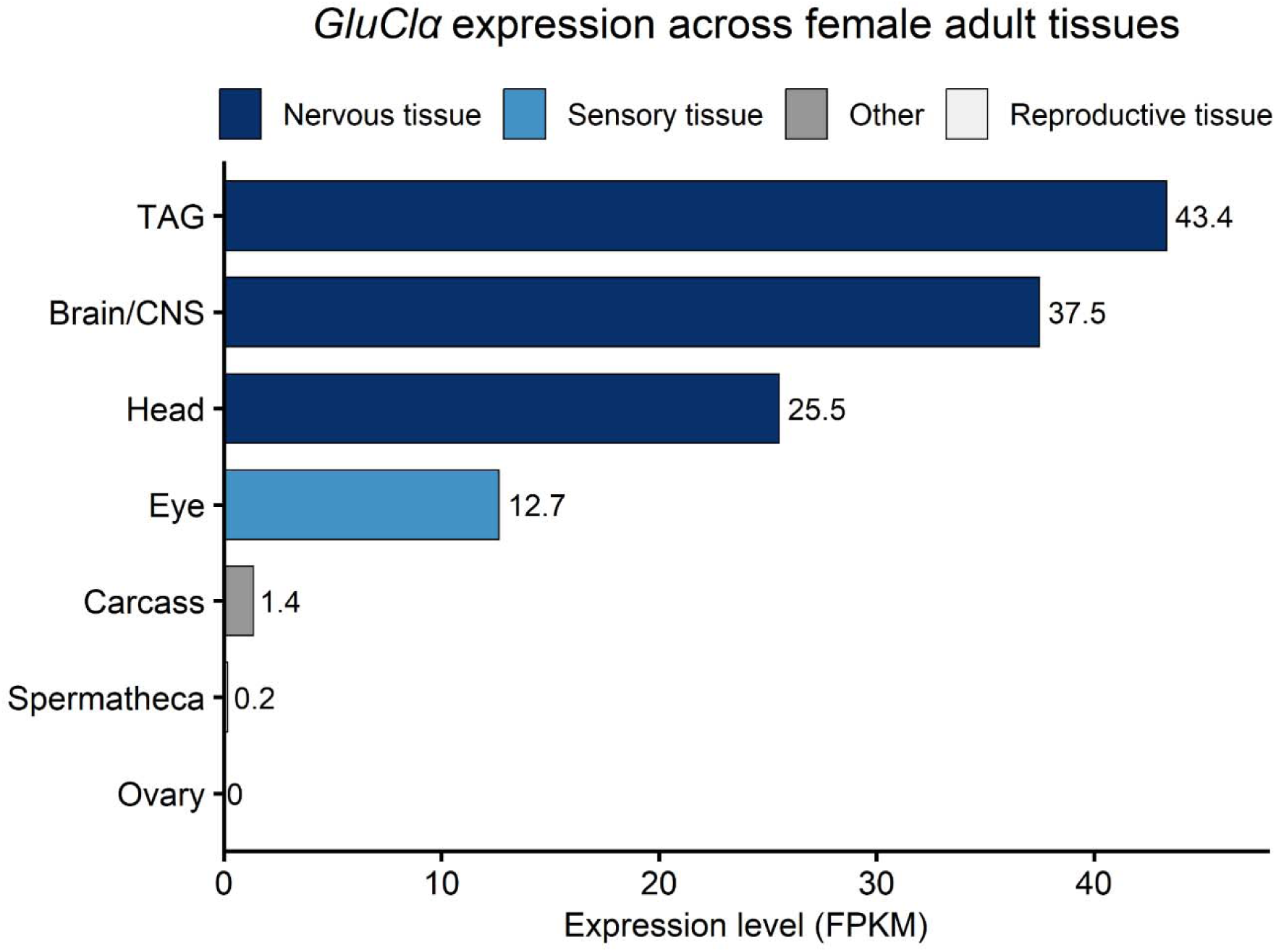
*GluCl*α is predominantly expressed in the nervous system of adult female *Drosophila*. Expression levels of *GluCl*α across adult female tissues obtained from the FlyAtlas2 database. Expression values are shown as fragments per kilobase of transcript per million mapped reads (FPKM).

**Fig. S7.**
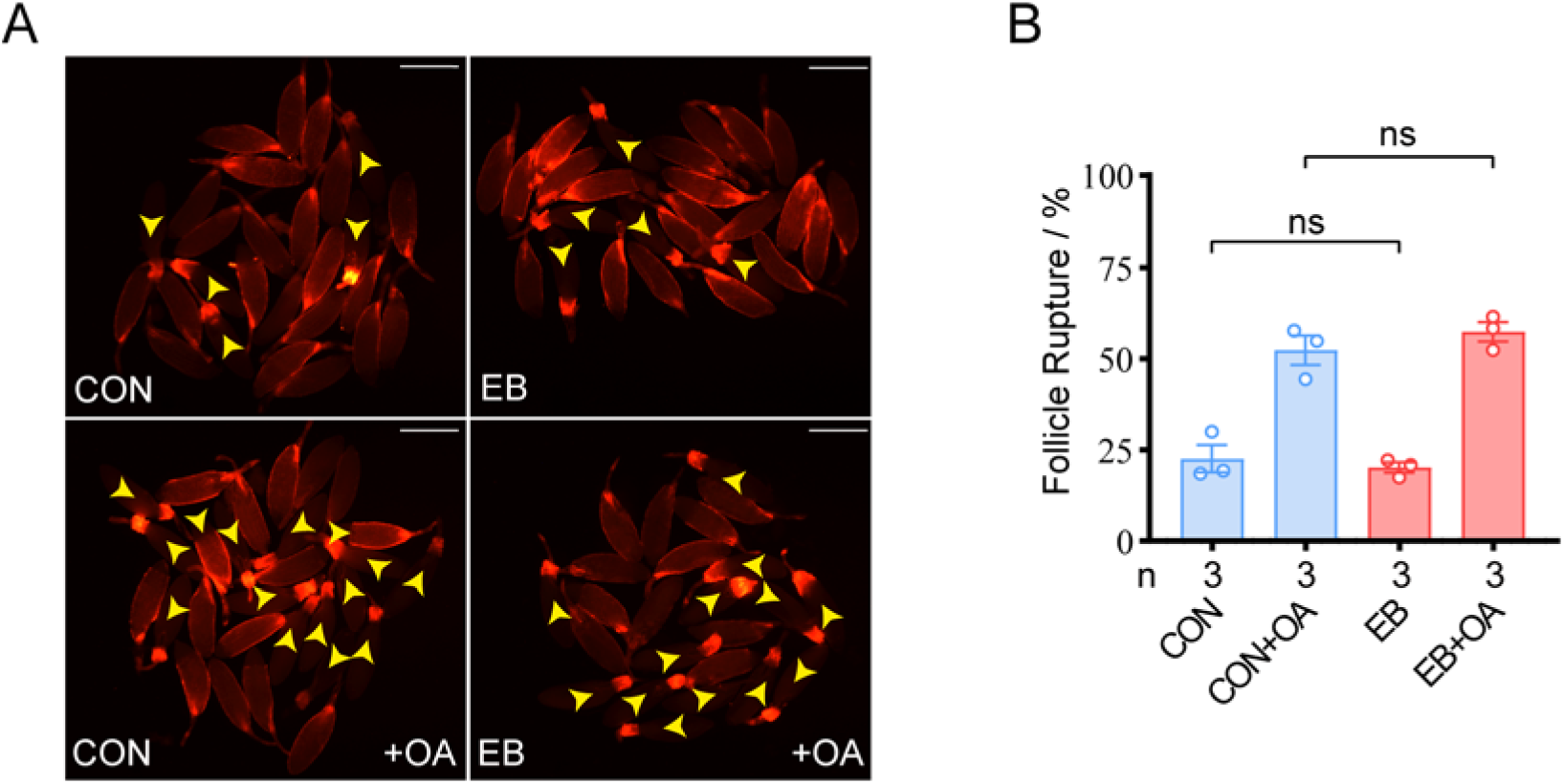
Follicle rupture following ex vivo treatment with EB and OA. (A) Representative images of follicle rupture following ex vivo treatment with EB, OA, or their combination. Arrowheads indicate ruptured follicles. Scale bars, 100 μm. (B) Quantification of follicle rupture shown in (A). Data are presented as mean ± s.e.m. Statistical significance was determined using one-way ANOVA followed by Tukey’s multiple comparison test. ns, not significant.

**Fig. S8.**
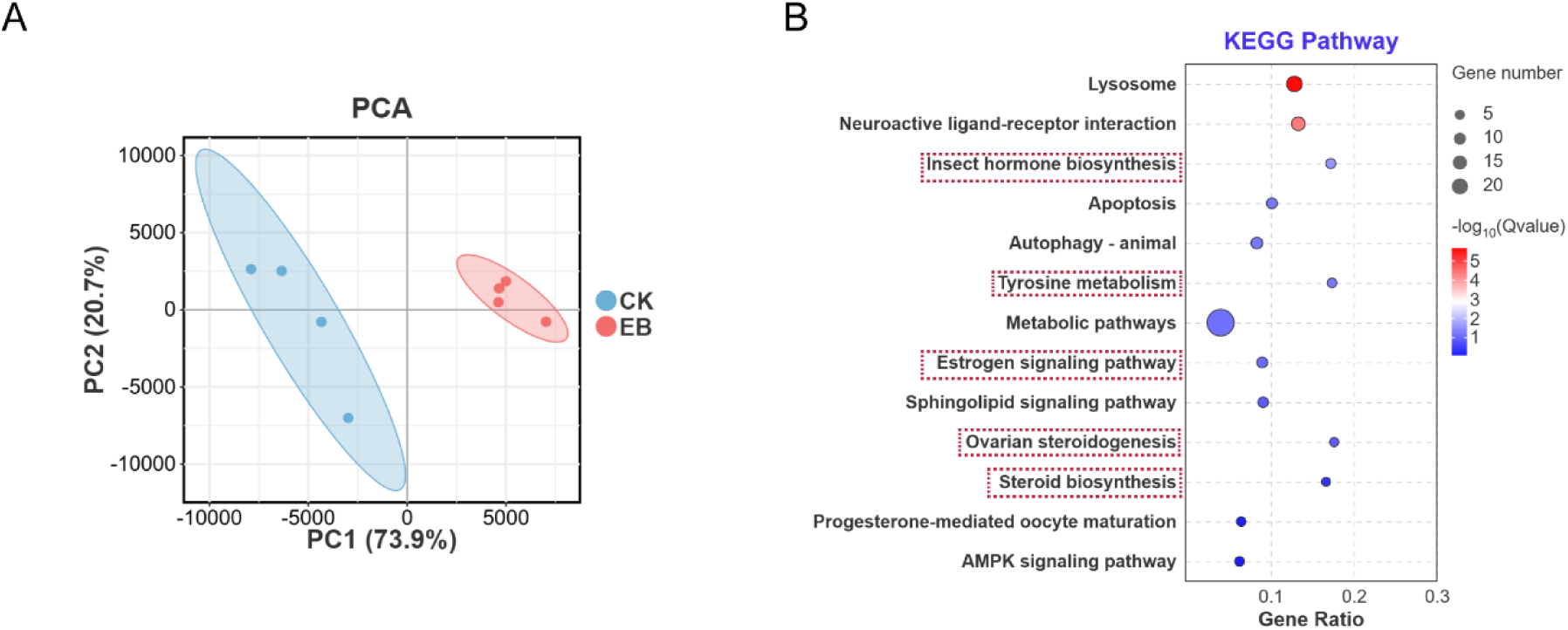
Global transcriptomic profiling of control and EB-treated females. (A) Principal component analysis (PCA) of transcriptomic datasets from control (CK) and EB females. Each dot represents an independent biological replicate. (B) KEGG pathway enrichment analysis of differentially expressed genes between control and EB-treated females. Dot size indicates gene number, and color represents −log10(Q value).

**Fig. S9.**
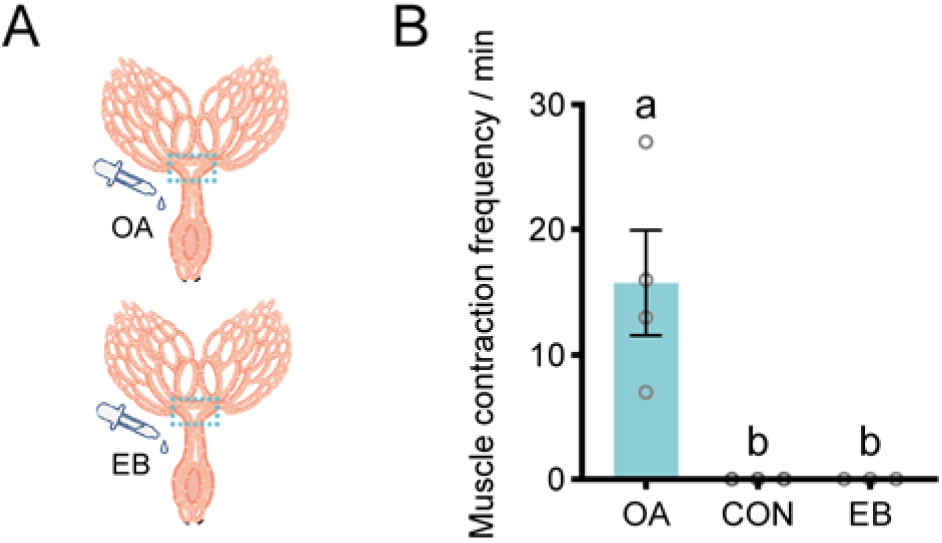
Ex vivo measurement of oviduct muscle contraction following octopamine and emamectin benzoate treatment. (A) Schematic illustration of the ex vivo assay used to measure oviduct muscle contraction following treatment with octopamine (OA) or emamectin benzoate (EB). (B) Quantification of oviduct muscle contraction frequency under the indicated treatment conditions. Data are presented as mean ± s.e.m. Statistical significance was determined using one-way ANOVA followed by Tukey’s multiple comparison test. Different letters indicate significant differences among groups (*P < 0.05*)

**Table S1.** Determination of the toxicity of Emamectin Benzoate to *Drosophila* by artificial diet incorporation method.

| Tested stages | Slope ± SE | LC<sub>50</sub> (95%F.L.)<br>(mg/L) | $\chi^2$(df) | P value |
| --- | --- | --- | --- | --- |
| Female | 4.968±0.649 | 31.971<br>(24.578-42.017) | 2.9468 (4) | 0.57 |
| Male | 3.103±0.387 | 30.762<br>(25.802-36.901) | 1.108 (4) | 0.89 |
| Larva | 3.001±0.352 | 0.00099<br>( 0.0008-0.0011 ) | 1.978 (4) | 0.74 |

